# Programming Brain Cell-Type-Selective Delivery In Vivo with Transporter-Guided Therapeutics

**DOI:** 10.64898/2026.06.14.732141

**Authors:** Roshan W. Gunasekara, Lejie Zhang, Lei Tong, John Zhou, Hoang Kim Trinh, Sean Pinon, Madison Gendreau, Emily Scott, John Chiari, Jaime Grutzendler

## Abstract

Many diseases arise from dysfunction of defined cell populations, yet most therapeutics distribute broadly, limiting efficacy and causing toxicity. We developed ExACT, a platform for cell-type-selective intracellular delivery that exploits membrane transporters. In vivo screening of combinatorial fluorescent small-molecule libraries in mouse brain identified chemistries whose uptake is dictated by endogenous transporter expression, yielding compounds with preferential entry into neurons, astrocytes, pericytes and endothelial cells. One series showed strong selectivity for brain and retinal endothelium, where Slco1a4 mediated uptake. This selectivity principle extended to the human orthologue SLCO1A2, highly expressed in brain endothelium and oligodendrocytes, where it mediated selective uptake in a humanized mouse model and human iPSC-derived oligodendrocytes. Ectopic expression of SLCO1A2 in neurons via gene therapy created a synthetic entry port, conferring ExACT conjugate uptake on otherwise inaccessible cells. Bifunctional compounds linking transporter-targeting motifs to antisense oligonucleotides or small-molecule drugs retained pharmacological activity while conferring transporter-dependent cell-type selectivity, illustrating how transporter diversity can be harnessed for precision pharmacotherapy.

## Introduction

The pathophysiology of many disorders involves mechanisms that selectively affect specific cell types within distinct organs, resulting in complex patterns of cellular vulnerability and dysfunction. In the central nervous system (CNS) and retina, certain cell types can exhibit selective vulnerability.^1–7^ This poses a challenge for drug development, as targeting common signaling pathways can have beneficial or harmful effects depending on the affected cell types and their roles within intricate tissue networks.^8^ To address these challenges, advances in cell-type selective pharmacotherapy, especially in the immune-oncology fields, using antibodies,^9^ peptides,^10^ and antibody-drug conjugates,^11^ have demonstrated the potential to target specific cell populations. Additionally, nanoparticles, liposomes, and viral vectors engineered with cell-specific ligands or capsids^12^ are enabling more precise delivery of therapeutics, improving efficacy and minimizing off-target effects.^13^

Despite notable advances in targeted delivery, achieving selective cellular targeting within the CNS remains a major challenge. This is especially true for small molecules, which remain a mainstay of pharmacotherapy, and emerging nucleic acid-based therapies, such as antisense oligonucleotides (ASOs) and small interfering RNA (siRNA). Once delivered either systemically or by intrathecal brain injection, neither platform has the ability to selectively target specific cell types, including neurons, astrocytes, or endothelium. Consequently, these therapeutics distribute broadly across neural tissue, affecting both disease-relevant and non-target cells.^14^

To address these limitations, it is essential to identify new strategies that can endow therapeutics with cell-type-selective precision. One promising avenue lies in the unique expression patterns of membrane transporters, which facilitate the movement of substances like amino acids, nucleosides, and hormones and are often selectively enriched in particular cell populations.^15, 16^ In the brain, studies have shown that certain small molecule fluorophores are preferentially internalized by specific cell types, including pericytes,^17^ neurons,^18^ astrocytes,^19^ and oligodendrocytes,^20^ likely due to their affinity for distinct membrane transporters present in these cells. Building on these observations, systematic identification of small molecules with high specificity for particular transporters offers a rational foundation for designing pharmacological agents that can selectively target defined cell types, thereby overcoming current barriers to precision in CNS therapeutics.

Here we developed ExACT, for Engineered Therapeutics with Active Cell-Type-Selective Transport, as a platform to harness endogenous membrane transporters for cell-type-selective intracellular delivery. Unlike antibody-drug conjugates, engineered nanoparticles, or capsid-based vectors, which require prior knowledge of cell-surface antigens or receptors, ExACT uses chemically defined uptake motifs to engage endogenous transporter activity and deliver linked therapeutic cargoes into selected cell populations. In this study, we combined *in vivo* chemical screening, transporter expression mapping, genetic perturbation and chemical diversification to identify transporter-harnessing motifs with cell-type-selective uptake in the CNS and retina. We then converted these motifs into bifunctional compounds capable of delivering pharmacological cargoes, including small molecule drugs and ASOs, with transporter-dependent selectivity. We further show that this strategy extends to human transporter orthologues and that ectopic transporter expression via viral vectors can expand targeting to cell types lacking endogenous transporter activity. Together, these findings establish membrane transporters as modular entry pathways for engineered bifunctional small molecule compounds and define ExACT as a strategy for cell-type-selective therapeutic delivery.

## Results

### Generation of a Combinatorial Library of Small Molecule Fluorophores

We hypothesized that novel small molecules capable of selectively entering specific brain cell types could be identified by screening a fluorescent compound library through optical imaging in the live mouse brain. To test this, we generated a chemical library of approximately 1,200 fluorescent compounds through combinatorial synthesis^21^ (Fig. 1a, Schemes S1-S6). These compounds incorporated a wide diversity of functional groups and fluorophore backbones,^22^ including cyanine, rhodamine, coumarin, fluorescein, and BODIPY, yielding fluorescent compounds ranging from 300 to 600 Da. Synthetic efficiency was maximized through short reaction routes and microwave-assisted chemistry.^23^ The exclusive use of non-fluorescent starting materials eliminated the need for purification steps, enabling direct imaging-based screening without interference from fluorescent reactants.

**Fig. 1:**
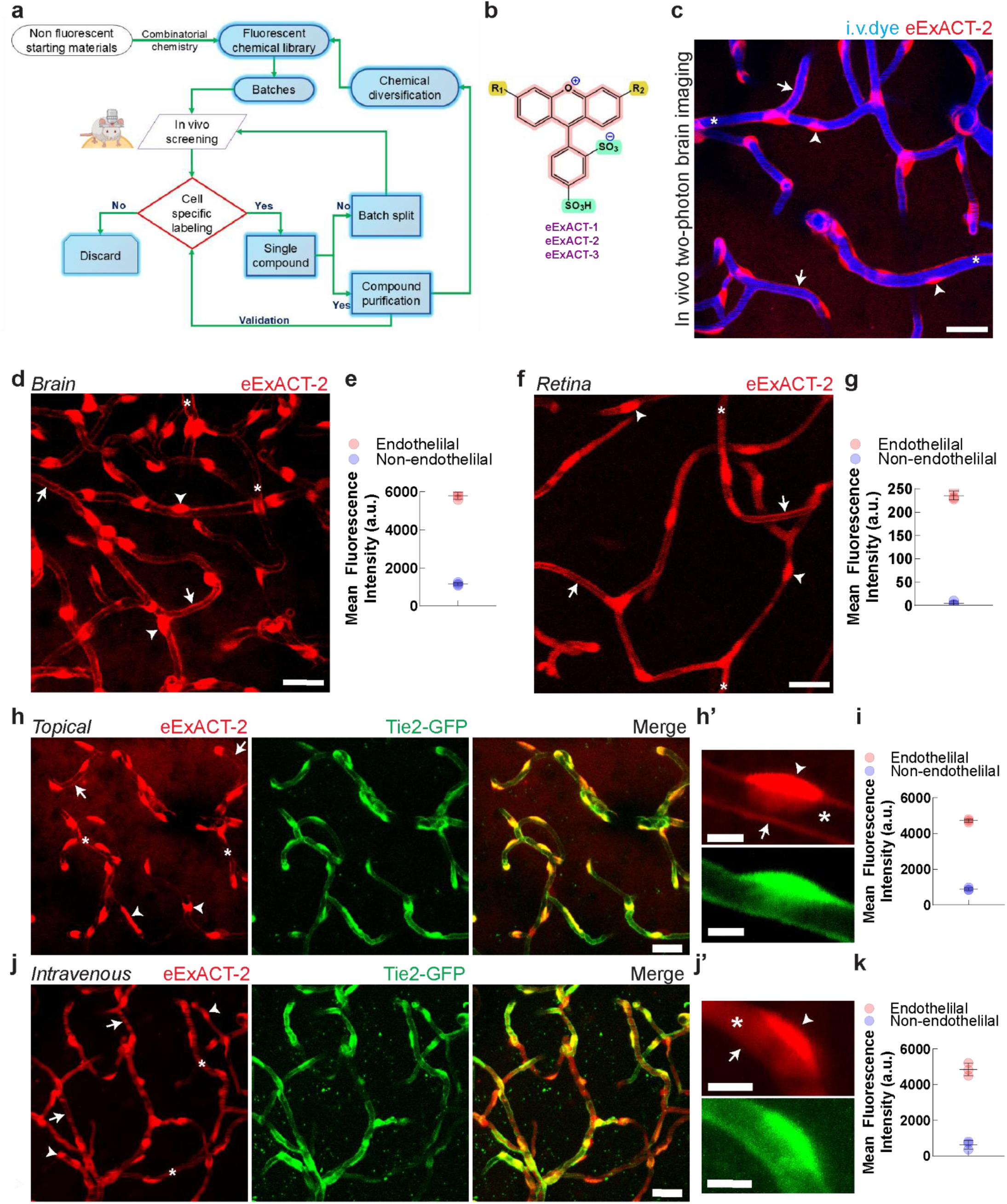
Intravital imaging–based screening of a combinatorial fluorophore library identifies endothelial-selective compounds (eExACT) in the brain and retina. **a.** Schematic overview of the screening strategy used to identify cell-specific fluorophores in the live mouse brain from a combinatorial chemical library. **b.** Chemical structures of three representative endothelial-specific compounds (eExACT-1, -2, and -3). **c.** Intravital two-photon image from the mouse cortex showing intact microvasculature after topical cortical application of eExACT-2 (50 µM, red) and intravascular blue dye injection (i.v. dye blue), highlighting normal intravascular perfusion and strong labeling of endothelial cell bodies (arrowheads), processes (arrows), and vessel lumen (asterisks). Scale bar, 20 μm. **d.** Intravital two-photon image from wild-type mouse cortex showing selective endothelial labeling following topical application of eExACT-2 (50 µM). Cell bodies (arrowheads), processes (arrows), and vessel lumen (asterisks). Scale bar, 20 μm. **e.** Quantification of mean fluorescence intensity in endothelial versus non-endothelial cells in the brain, 30 minutes after eExACT-2 cortical application (related to d). **f.** Confocal image of a mouse retina explant following *in vivo* intravitreal injection of eExACT-2 (50 µM), showing selective labeling of endothelial cells. Cell bodies (arrowheads), processes (arrows), and vessel lumen (asterisks). Scale bar, 20 μm. **g.** Quantification of mean fluorescence intensity in endothelial versus non-endothelial structures in the retina 30 minutes after eExACT-2 intravitreal injection (related to f). **h.** Two-photon imaging in Tie2-GFP reporter mice following cortical topical administration of eExACT-2 (50 µM) in the live mouse brain 30 minutes after cortical application, demonstrating colocalization between eExACT-2 fluorescence (red) and GFP-labeled endothelium (green). Cell bodies (arrowheads), processes (arrows), and vessel lumen (asterisks). Scale bar, 20 μm. **h’.** High-magnification view showing colocalization between GFP (green) and eExACT-2 (red) signals. Cell bodies (arrowheads); processes (arrows), and vessel lumen (asterisks). Scale bar, 5 μm. **i.** Quantification of mean fluorescence intensity in endothelial versus non-endothelial cells 30 minutes after eExACT-2 cortical application (related to h–h’). **j.** Two-photon *in vivo* imaging in Tie2-GFP mice following intravenous administration of eExACT-2 (100 µM). GFP (green) and eExACT-2 (red) colocalization is observed at endothelial cell bodies. Cell bodies (arrowheads); processes (arrows) and vessel lumen (asterisks). Scale bar, 20 μm. **j’.** High-magnification view showing precise colocalization between GFP and eExACT-2 signals. Fine endothelial processes are obscured due to the bright intravascular labeling. Cell bodies (arrowheads); processes (arrows) and vessel lumen (asterisks). Scale bar, 5 μm. **k.** Quantification of endothelial versus non-endothelial fluorescence 240 minutes after eExACT-2 intravenous delivery (related to j and j’).

### Library Screening Through Optical Imaging in the Live Mouse Brain

We reasoned that the most effective way to discover compounds with selective affinity for defined brain cell types and transport mechanisms is to screen them directly in the intact *in vivo* brain microenvironment. To do this, we developed a cranial window–based approach in which candidate molecules are applied topically to the cortical surface. This configuration uses substantially less material than intravenous dosing and avoids confounds from variable blood–brain barrier (BBB) permeability.

To increase throughput, we implemented a pooling strategy in which batches of ten unpurified compounds were dissolved at micromolar concentrations in artificial cerebrospinal fluid (ACSF) containing 3 percent (v/v) dimethyl sulfoxide (DMSO). These mixtures were applied topically to the exposed brain surface. We then used high-resolution intravital two-photon imaging to visualize fluorophore distribution, enabling precise localization of compound uptake and cellular targeting within the intact brain microenvironment (Fig. 1a).

This *in vivo* screen revealed a spectrum of labeling patterns. Most compound pools produced diffuse interstitial labeling consistent with nonspecific distribution, whereas a small subset showed striking cell-type specificity. To refine hit selection, pools with cell-specific labeling were subdivided and re-screened iteratively until individual compounds with distinct cellular uptake patterns were isolated. Because these initial hits were unpurified, we considered the possibility that the labeling arose from fluorescent byproducts. To address this, we scaled up synthesis of promising hits, purified them chromatographically, and repeated the *in vivo* assays to confirm their cell-type specificity (Fig. 1a). Through this iterative process, which involved an intravital screen of approximately 1,200 molecules (Schemes S1-S6), we identified ten compounds that exhibited selective uptake in defined cell types, including neurons, pericytes, astrocytes, and endothelial cells (Supplementary Figs. 1a–1f and Fig. 1b–1d). The recovery of only a small number of cell-selective compounds from this large library underscores the highly specific molecular interactions required for cellular uptake in the intact brain.

### Discovery of Compounds with Brain and Retina Endothelial Specificity

Following the initial screen, we carried out proof-of-concept studies on a subset of compounds that showed rapid uptake and exceptional selectivity for endothelial cells in both the brain and retina, which we termed endothelial ExACT (eExACT) (Figs. 1b–1g, Supplementary Figs. 1g–1j, and Supplementary Figs. 2a–2e). This specificity was evident from the characteristic morphology of labeled endothelial cell bodies and processes, as well as precise colocalization with endothelial cells in Tie2-GFP reporter mice (Figs. 1h–1i, Supplementary Figs. 1g–1j). To probe the underlying transport mechanism, we applied eExACT both topically, to expose the abluminal endothelial surface (Figs. 1h–1i, Supplementary Figs. 1g–1j), and intravascularly, to target the luminal surface (Figs. 1j–1k). Even at high concentrations, eExACT did not label non-endothelial cells in the cortex, underscoring its specificity (Figs. 1h–1k, Supplementary Figs. 2a–2e). The compounds selectively entered endothelial cells across arterioles, capillaries, and venules within the brain parenchyma (Figs. 1c–1e), and application of eExACT to acute brain slices confirmed robust endothelial uptake in cortical, subcortical, and white matter regions (Table S1). Likewise, intravitreal injection led to rapid and selective labeling of endothelial cells in retinal arterioles, venules, and capillaries (Figs. 1f–1g). Together, these experiments show that eExACT compounds possess high specificity and strong affinity for endothelial cells in both the brain and retina.

**Fig. 2:**
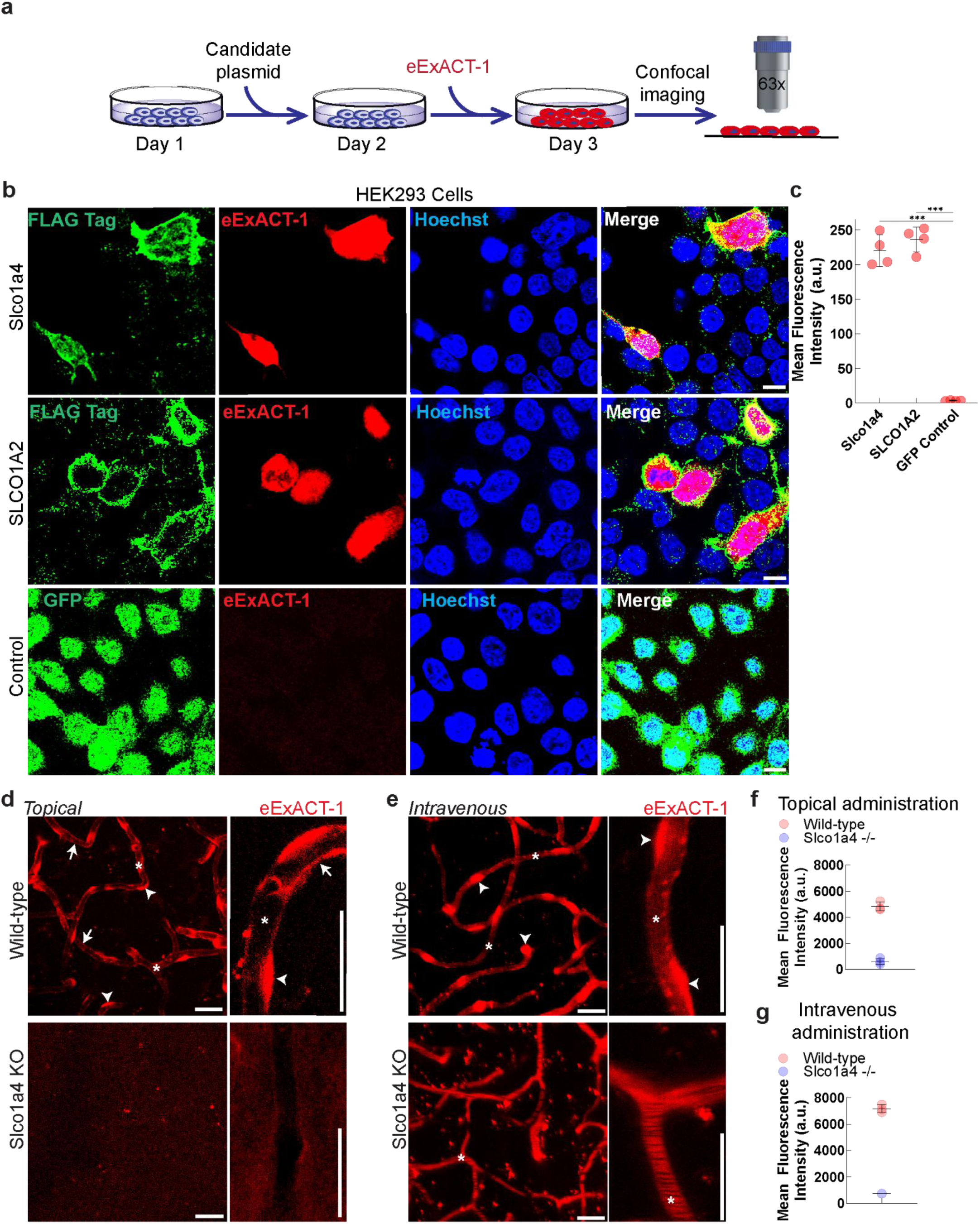
The solute carriers Slco1a4 (mouse) and SLCO1A2 (human) mediate selective intracellular uptake of eExACT. **a.** Schematic overview of the experimental workflow used for solute carrier transfections in HEK293 cells and confocal imaging protocol. **b.** Confocal images of HEK293 cells transfected with the solute carriers *Slco1a4* (mouse), *SLCO1A2* (human), or GFP control, following exposure to eExACT-1 (10 μM). Robust intracellular uptake is observed only in cells expressing either solute carrier. The *Slco1a4* and *SLCO1A2* proteins were fused to a FLAG tag for visualization. Scale bar, 10 μm. **c.** Quantification of mean fluorescence intensity in HEK293 cells transfected with the solute carriers *Slco1a4* (mouse), *SLCO1A2* (human), or GFP control, after eExACT-1 exposure (related to b). *** indicates P<0.003 (unpaired t-test). **d.** *In vivo* two-photon images of the cortex from wild-type mice (top) and Slco1a4 deficient mice (bottom) after topical application of eExACT-1 (50 μM, red). In wild-type mice, eExACT-1 labels endothelial processes (arrows) and cell bodies (arrowheads). In contrast, signal is absent in Slco1a4 deficient mice, confirming transporter dependence. Vessel lumen (asterisks). Scale bars, 20 μm. **e.** *In vivo* two-photon images of the cortex following intravenous administration of eExACT-1 (100 μM, red) in wild-type (top) and Slco1a4 deficient mice (bottom). Wild-type mice show uptake in endothelial cell bodies (arrowheads), whereas knockout mice lack uptake despite visible intravascular eExACT-1 fluorescence (asterisks). Scale bar, 20 μm. **f.** Quantification of mean fluorescence intensity in endothelial cells of wild-type versus Slco1a4 deficient mice, 30 minutes after eExACT-1 cortical application (related to d). **g.** Quantification of mean fluorescence intensity in endothelial cells of wild-type versus Slco1a4 deficient mice, 240 minutes after eExACT-1 intravenous application (related to e).

### Mechanisms of Endothelial Uptake and Cell-Type Specificity

We next investigated the mechanisms underlying cellular uptake of eExACT. Given their endothelial selectivity, relatively high molecular weight and hydrophilicity (450–650 Da, Fig. 1b), we hypothesized that uptake is mediated by specific membrane transporters. We further postulated that the cell-type specificity of eExACT reflects differential expression of these transporters in brain endothelial cells compared to other cell types. To explore this, we analyzed publicly available single-cell RNA-sequencing datasets from mouse brain,^24, 25^ and identified seven solute carrier (SLC) transporters with endothelial-enriched expression as candidate mediators of eExACT uptake (Table S2 and Supplementary Figs. 3a–3g).

**Fig. 3:**
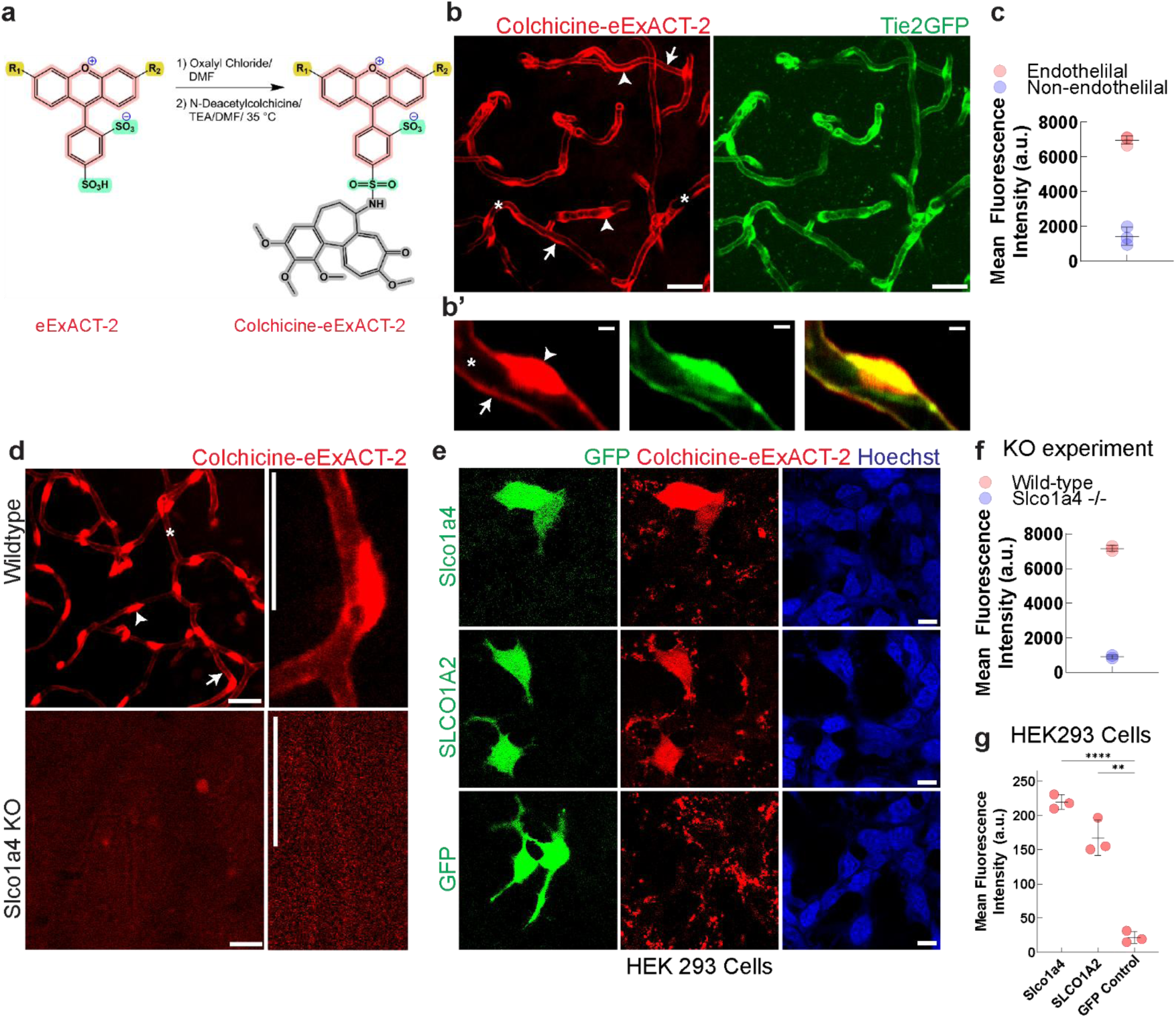
Conjugation of eExACT to a pharmacological agent preserves solute carrier–dependent cellular uptake. **a.** Schematic of the synthetic route used to conjugate Colchicine to eExACT-2**. b.** Two-photon imaging in Tie2-GFP reporter mice following cortical topical administration of Colchicine-eExACT-2 (50 µM) in the live mouse brain 30 minutes after cortical application, demonstrating colocalization between Colchicine-eExACT-2 fluorescence (red) and GFP-labeled endothelium (green). Cell bodies (arrowheads), processes (arrows), and vessel lumen (asterisks). Scale bar, 20 μm. **b’.** High-magnification view showing colocalization between GFP (green) and Colchicine-eExACT-2 (red) signals. Cell bodies (arrowheads), processes (arrows), and vessel lumen (asterisks). Scale bar, 5 μm. **c.** Quantification of mean fluorescence intensity in endothelial versus non-endothelial cells 30 minutes after Colchicine-eExACT-2 cortical application (related to b–b’). **d.** *In vivo* two-photon images of the cortex following topical administration of Colchicine-eExACT-2 (50 μM) in wild-type (top) and *Slco1a4* deficient (bottom) mice. Uptake in endothelial processes (arrows) and cell bodies (arrowheads) is eliminated in knockout mice. Vessel lumen (asterisks). Scale bar, 20 μm. **e.** Confocal images of HEK293 cells transfected with either mouse *Slco1a4*, human *SLCO1A2*, or GFP control, following administration of Colchicine-eExACT-2 (10 μM). Robust and selective uptake is observed in *Slco1a4* and *SLCO1A2*-expressing cells but not in GFP controls. Scale bars, 10 μm. **f.** Quantification of mean fluorescence intensity in endothelial cells of wild-type versus slco1a4 deficient mice, 30 minutes after Colchicine-eExACT-2 cortical application (related to d). **g.** Quantification of mean fluorescence intensity in HEK293 cells transfected with the solute carriers *Slco1a4* (mouse), *SLCO1A2* (human), or GFP control, after Colchicine-eExACT-2 exposure (related to e). ** indicates P<0.01 and **** indicates P<0.0001 (unpaired t-test).

To test these candidates, we individually overexpressed each transporter in human embryonic kidney (HEK 293) cells,^26^ which do not take up eExACT under baseline conditions (Fig. 2a and Supplementary Figs. 4a–4b). Only overexpression of the mouse organic anion transporter Slco1a4 produced a marked increase in intracellular eExACT accumulation (Figs. 2b and 2c, Supplementary Figs. 3a–3g, Supplementary Figs. 4a–4b, and Table S2). We then examined SLCO1A2, the human ortholog of Slco1a4,^27, 28^ and found that its overexpression also conferred robust eExACT uptake in HEK293 cells (Figs. 2b–2c), indicating a conserved transport mechanism between mouse and human.

**Fig. 4:**
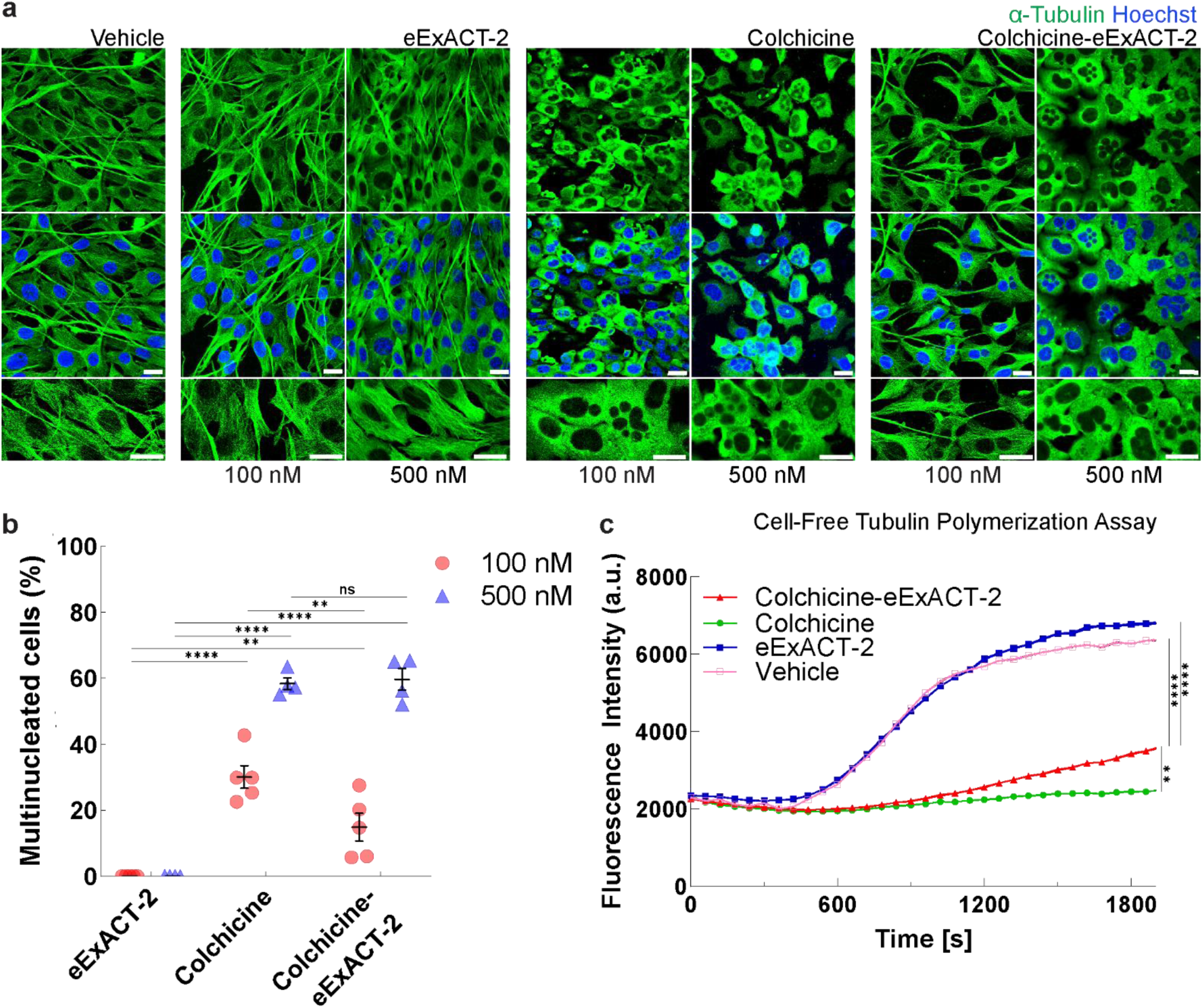
eExACT-drug conjugate retains the bioactivity of linked pharmacological compound. **a.** Confocal images of α-tubulin immunofluorescence (green) in NIH3T3 cells treated with vehicle, eExACT-2, Colchicine, or Colchicine–eExACT-2 at two concentrations (100 nM and 500 nM). The normal filamentous microtubule pattern seen in vehicle- and eExACT-treated cells is disrupted following treatment with Colchicine or Colchicine–eExACT-2, resulting in diffuse tubulin staining (green), altered cell morphology, and increased multinucleation (Hoechst, blue). Scale bars, 20 μm. **b.** Quantification of multinucleated cells across treatments and concentrations (N = 3–5 wells per group, with 3–5 images analyzed per well; total cells per group >50). ** indicates P<0.01, **** indicates P<0.0001, and ns indicates not significant (unpaired t-test). **c.** Fluorescence-based cell-free tubulin polymerization assay comparing the effects of Vehicle, eExACT-2, Colchicine–eExACT-2, and Colchicine (3 µM). Curves show mean fluorescence intensity over time (N = 3 technical replicates per group). For statistical analysis of panel C, Two-sample Kolmogorov-Smirnov test was used to assess overall differences between groups where ** indicates P=0.013 and **** indicates P<0.0001.

To confirm the role of Slco1a4 *in vivo*, we used mice with a deletion of the Slco1a/1b gene cluster, which includes Slco1a4, the only member highly expressed in brain endothelium (Supplementary Figs. 5a–5e). Topical application of eExACT to the cortex (Figs. 2d and 2f, Supplementary Fig. 6a) or intravenous injection (Figs. 2e and 2g), followed by intravital two-photon imaging, revealed a complete loss of endothelial labeling in these mice, strongly implicating Slco1a4 as essential for eExACT uptake *in vivo*. To assess regional differences in uptake, we applied eExACT topically and imaged both intraparenchymal microvessels and extraparenchymal leptomeningeal arteries. This revealed striking spatial segregation, with intraparenchymal vessels showing robust labeling, whereas larger leptomeningeal arteries exhibited minimal to no uptake (Supplementary Fig. 7a), consistent with Slco1a4 expression being largely confined to the intraparenchymal microvasculature.

**Fig. 5:**
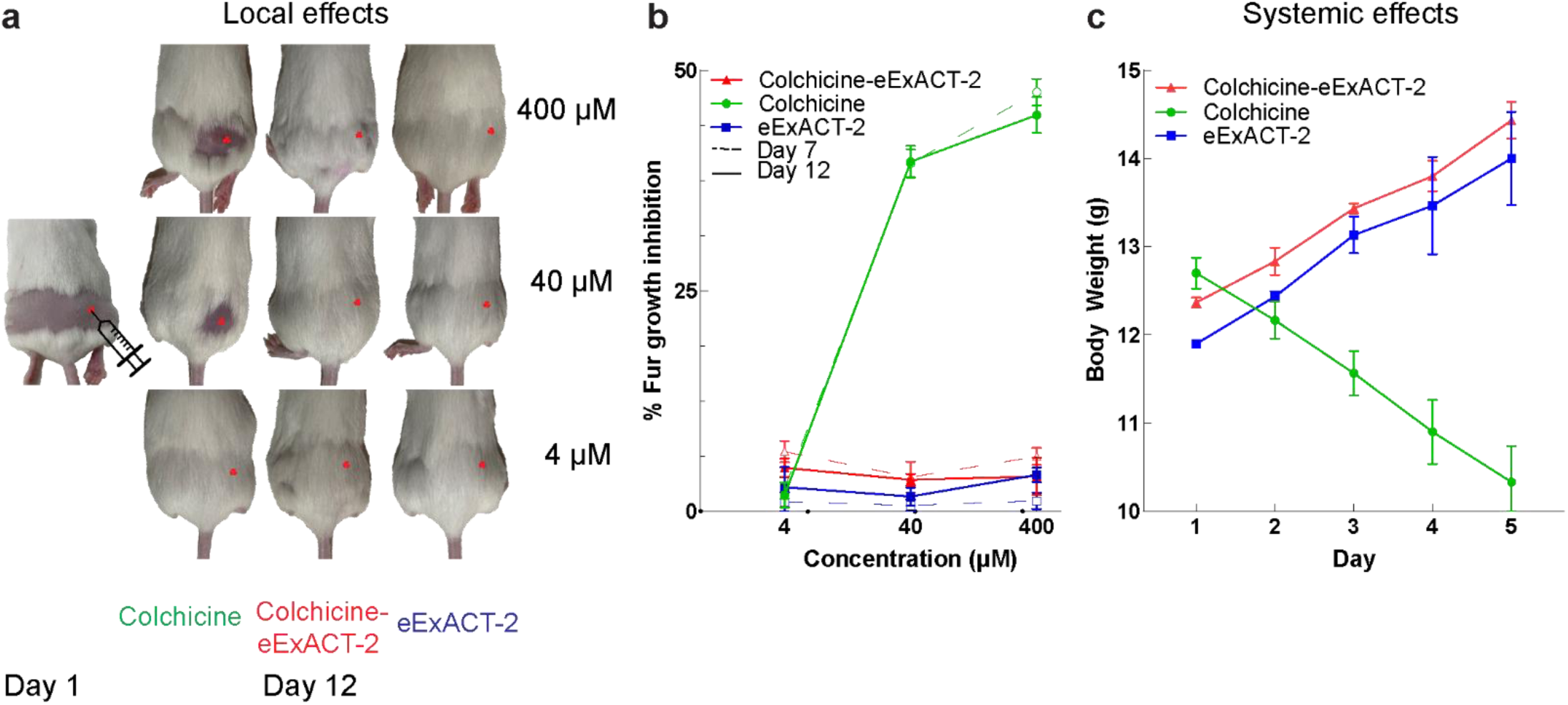
Conjugation of Colchicine to eExACT-2 reduces local and systemic toxicity. **a.** To assess local colchicine toxicity, fur was shaved from the lower back of mice, and treatments (4 µM, 40 µM, or 400 µM) were administered intradermally in the right quadrant (red circle), while the left quadrant served as an internal control. Treatments included Colchicine (green), Colchicine–eExACT-2 (red), and eExACT-2 (blue). Hair regrowth was monitored and quantified over time across different treatment groups and doses. **b.** Quantification of fur regrowth by measuring grayscale intensity differences between injected and uninjected quadrants, reflecting compound-specific effects on follicular stem cell proliferation. **c.** Body weight monitoring in 3-week-old mice receiving daily intraperitoneal injections of Colchicine, Colchicine–eExACT-2, or eExACT-2 (2.5 µmol/g) for five consecutive days. Mean ± SEM, N = 3 mice per group.

### Development of Drug Conjugates with Cell-Type Specific Intracellular Uptake

Having identified the membrane transport mechanism for eExACT, we hypothesized that these compounds could be conjugated to pharmacological agents while preserving endothelial cell specificity. Our first objective was to define chemical positions on eExACT that tolerate conjugation without disrupting transporter-mediated uptake. We therefore devised a strategy to map functional groups essential for cell-type-specific transport and to selectively modify noncritical positions. This iterative process combined systematic diversification of functional groups on the eExACT scaffold (Supplementary Figs. 8a–8c and 9a–9c) with *in vivo* two-photon brain imaging to assess uptake, enabling direct structure–property relationship (SPR) analysis in the intact brain. Using this approach, we synthesized and screened approximately 20 modified eExACT derivatives. From this series, we identified a sulfonate group (–SO₃H) as dispensable for endothelial specificity, making it an ideal handle for derivatization through conjugation with amines. Leveraging this handle, we generated a panel of eExACT–drug conjugates by coupling amine-containing small molecules, including FDA-approved drugs with molecular weights up to ∼1.2 kDa (Supplementary Figs. 9a–9c). Notably, these conjugates retained the rapid endothelial uptake and cellular specificity of the parent compound, despite increased hydrophilicity and size (Supplementary Figs. 9a–9c), demonstrating the feasibility of eExACT as a modular platform for cell-type-specific drug delivery.

To further evaluate this conjugation strategy, we selected colchicine, a well-characterized drug with established chemistries that preserve its pharmacological activity.^29–31^ Colchicine itself lacks intrinsic cell-type-specific uptake but has robust, quantifiable effects through microtubule inhibition and anti-inflammatory activity, making it a useful model compound for assessing target engagement.^32, 33^ For conjugation, we first converted eExACT into a sulfonyl chloride intermediate via reaction with oxalyl chloride, then coupled this intermediate to N-deacetylcolchicine to generate a bifunctional colchicine–eExACT conjugate (Fig. 3a).

To assess *in vivo* targeting, we applied the colchicine–eExACT conjugate topically to the brain and performed two-photon imaging, which confirmed selective uptake by endothelial cells (Figs. 3b–3c). In mice lacking Slco1a4, endothelial uptake was completely abolished (Figs. 3d and 3f, Supplementary Fig. 6b), further validating the requirement of this transporter for intracellular delivery *in vivo*. To determine whether the conjugate retained the uptake specificity of the unconjugated parent compound, we examined its uptake in HEK293 cells transfected with Slco1a4 or SLCO1A2. In both cases, we observed efficient intracellular accumulation (Figs. 3e and 3g), indicating that the colchicine–eExACT conjugate preserves Slco1a4/SLCO1A2-dependent cell-type-specific transport.^34^

### Drug Conjugates Preserve Pharmacological Properties While Markedly Reducing Systemic Toxicity

After confirming that colchicine–eExACT retained selective endothelial uptake, we next asked whether the conjugate preserved colchicine’s pharmacological activity. We used NIH 3T3 fibroblasts, a well-established model for studying microtubule polymerization,^35^ which also take up eExACT at baseline. High-resolution tubulin immunofluorescence revealed a dose-dependent disruption of microtubule architecture following colchicine–eExACT treatment (Figs. 4a –4b), and at nanomolar concentrations cells exhibited multinucleation, consistent with colchicine’s antimitotic effects.^36^ These findings were corroborated in a cell-free microtubule polymerization assay^32^, in which the colchicine–eExACT conjugate caused robust, dose-dependent inhibition of polymerization (Fig. 4c).

To compare toxicity of colchicine and its eExACT-conjugated form, we performed *in vivo* studies in mice. Colchicine is known to cause systemic adverse effects, including bone marrow suppression and inhibition of hair growth.^37–39^ Prior work has shown that subcutaneous colchicine injections inhibit hair follicle stem-cell division and prevent fur regrowth after shaving.^37^ We reproduced this model, administering high concentrations (up to 400 μM) of either colchicine or colchicine–eExACT. As expected, colchicine blocked fur regrowth, whereas colchicine–eExACT permitted normal hair regrowth at equivalent dosing (Figs. 5a –5b). We further assessed systemic toxicity by monitoring weight gain in developing (P10) mice treated intraperitoneally for five days. Colchicine treatment caused significant weight loss, whereas colchicine–eExACT and unconjugated eExACT had no effect on normal growth (Fig. 5c), indicating a markedly reduced toxicity profile for the conjugate.^40^

To investigate the basis of this reduced toxicity, we examined tissue biodistribution of colchicine–eExACT using its intrinsic fluorescence. Following intravenous or topical administration of a high dose, we observed no appreciable uptake in endothelial cells of peripheral organs, in contrast to the robust labeling in brain and retina (Figs. 1d–1g and 3b–3c). In particular, heart, spleen, and skeletal muscle showed neither endothelial nor parenchymal uptake (Supplementary Figs. 10a–10d and 10g, Supplementary Figs. 11a–11c and 11f), consistent with limited Slco1a4 expression in these tissues.^41^ Notable exceptions were kidney and liver, which displayed strong uptake in renal tubular epithelium and hepatocytes, respectively, despite minimal Slco1a4 expression^41^ and absence of endothelial labeling by colchicine–eExACT (Supplementary Figs. 10e–10g and 11d–11f). This pattern likely reflects the central roles of these organs in small-molecule metabolism and clearance rather than Slco1a4-dependent endothelial transport. To further probe hepatic uptake mechanisms, we tested whether other solute carrier transporters expressed in hepatocytes, SLCO1B1, SLCO1B3, or SLCO2B1,^42^ could mediate eExACT uptake; overexpression of these transporters in HEK293 cells did not confer detectable uptake (Supplementary Figs. 12a – 12b), suggesting that liver accumulation occurs through alternative pathways.

Taken together, these findings indicate that the reduced systemic toxicity of colchicine–eExACT primarily reflects the restricted expression of Slco1a4 and the limited uptake of the larger, hydrophilic conjugate in most tissues. Importantly, despite minimal off-target distribution, the conjugate maintains full pharmacological activity at the intended site, supporting its potential as a cell-specific therapeutic platform with markedly reduced systemic side effects.

### eExACT Enables Targeted Delivery of Large Therapeutic Cargo Such as Antisense Oligonucleotides

We initially explored the upper size limit of deliverable cargo by conjugating eExACT to large molecules, including Oligonucleotide sequences of up to 6 kDa. Unexpectedly, two-photon imaging revealed rapid and robust endothelial uptake of the Oligo6k-eExACT conjugate *in vivo* (Supplementary Figs. 13a–13c), suggesting that even large, hydrophilic cargoes could be effectively internalized.

Building on these findings, we next tested whether eExACT could be extended to deliver nucleic acid–based therapeutics. We selected Malat1, a long non-coding RNA that is ubiquitously expressed and widely used for evaluating ASO delivery.^43^ To evaluate delivery, we conjugated an antisense oligonucleotide (ASO) targeting Malat1 to an eExACT scaffold and assessed its uptake *in vitro* and *in vivo* (Table S3). Two-photon imaging in live mice revealed robust and selective uptake of the Malat1–eExACT ASO conjugate in brain endothelial cells (Figs. 6a – 6b), consistent with transporter-mediated delivery. In contrast, a Malat1 ASO labeled with a standard Cy5 fluorophore showed no detectable uptake (Figs. 6c – 6d), underscoring the essential role of the eExACT scaffold for cell-specific entry. Additionally, when the conjugate was administered to Slco1a4-deficient mice, no endothelial uptake was observed (Figs. 6e –6f), providing further *in vivo* evidence that eExACT-mediated delivery depends on this transporter. Supporting these findings, uptake of the Malat1–eExACT ASO conjugate in HEK293 cells was strictly dependent on overexpression of SLCO1A2, the human ortholog of mouse Slco1a4 (Figs. 6g – 6h), confirming the requirement for transporter expression in vitro.

**Fig. 6:**
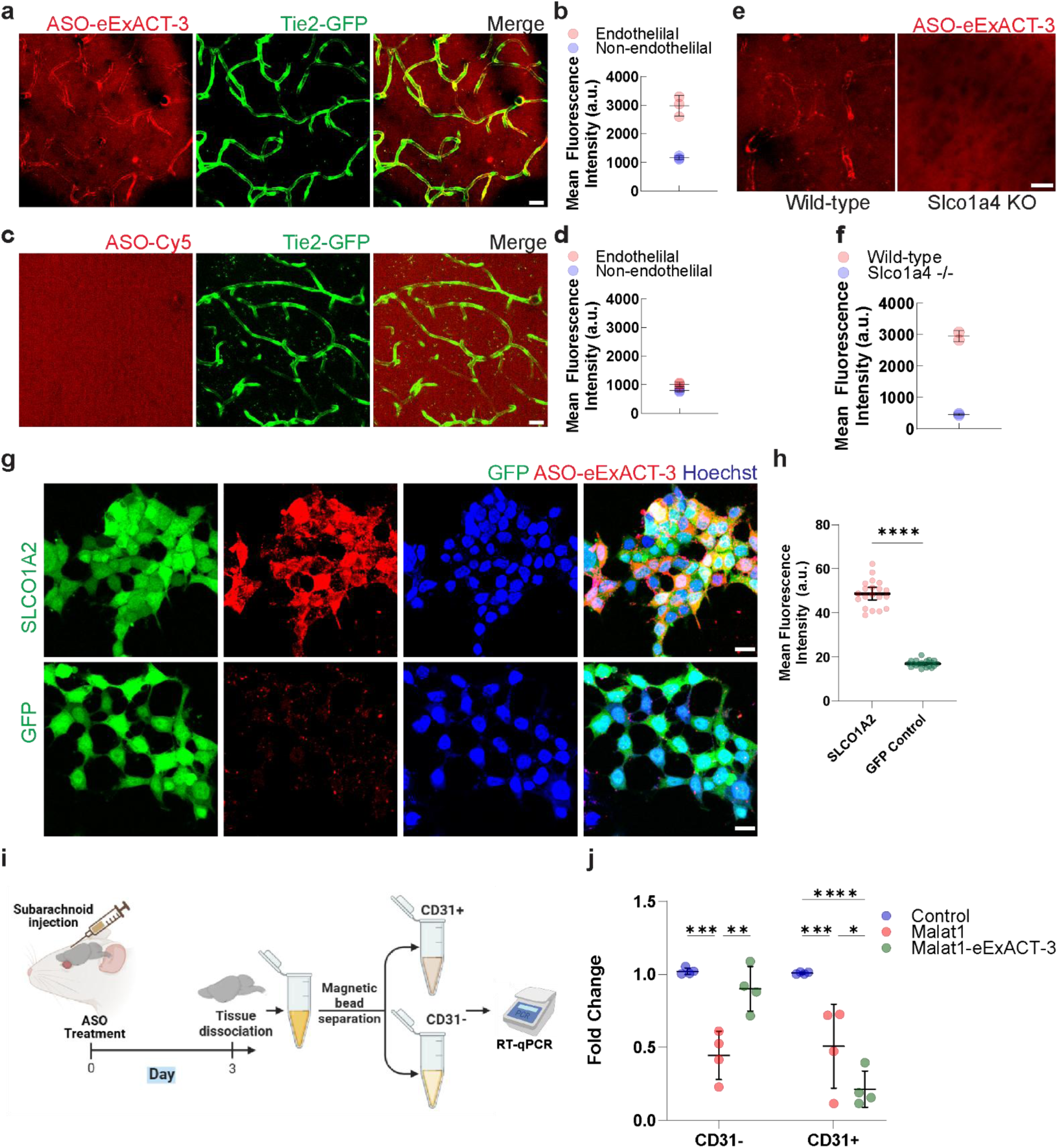
eExACT enables targeted delivery of large molecular cargo such as antisense oligonucleotides by promoting transporter-dependent cellular uptake and achieving gene silencing in vivo. **a, c.** *In vivo* two-photon imaging of Tie2-GFP mouse cortex 20 minutes after topical administration of ASO-eExACT-3 (a) or control ASO-Cy5 (c) (80 μM), shows selective uptake of the eExACT-conjugated ASO by brain endothelial cells. Scale bars, 20 μm. **b, d.** Quantification of fluorescence signal intensity in endothelial versus non-endothelial cells 20 minutes after topical application confirms selective uptake of ASO-eExACT-3 (**b**, 80 μM), which is absent with control ASO-Cy5 (**d**, 80 μM). **e–f.** Two-photon imaging (e) and quantification (f) 20 minutes after topical application show that ASO-eExACT (80 μM) delivery is abolished in Slco1a4 knockout mice compared to wild-type controls, indicating dependence on the Slco1a4 solute carrier. Scale bars, 20 μm. **g–h.** Confocal microscopy of HEK293 cells transfected with SLCO1A2 or GFP control (green) and treated with ASO-eExACT-3 (red, 1 μM) shows robust uptake in SLCO1A2-expressing cells (g), which is absent in control cells. Nuclei were counterstained with Hoechst (blue). Scale bars, 20 μm. Quantification of fluorescence intensity (h) confirms this difference (****p < 0.0001, unpaired t-test). **i.** Schematic overview of the experimental workflow. Mice received subarachnoid injections of either ASO-eExACT-3, ASO, or Control ASO (2.5 nmol), followed by brain extraction and dissociation into single-cell suspensions. Magnetic bead-based separation using anti-CD31 antibodies was then used to isolate endothelial (CD31⁺) and non-endothelial (CD31⁻) cell populations for downstream RNA analysis. **j.** RT-qPCR analysis of *Malat1* RNA expression in CD31⁻ and CD31⁺ cell fractions isolated 3 days after subarachnoid injection of *Malat1*-targeting ASO-eExACT-3 (*Malat1*-eExACT-3), unconjugated *Malat1* ASO, or control ASO (2.5 nmol). A significant reduction in *Malat1* expression was observed in the endothelial-enriched CD31⁺ fraction specifically in the *Malat1*-eExACT-3 treated group, indicating effective *in vivo* gene silencing and transporter-mediated target engagement. In contrast, the CD31⁻ fraction showed minimal *Malat1* knockdown with *Malat1*-eExACT treatment, unlike the broad effect of unconjugated *Malat1* ASO, highlighting enhanced cell-type specificity conferred by the eExACT platform. Bars represent mean ± SEM; N = 4 mice/group. Statistical comparisons were performed using two-way ANOVA with Tukey’s multiple comparisons test.

To evaluate target engagement, we delivered the Malat1–eExACT ASO *in vivo* via subarachnoid injection (Fig. 6i) and observed a robust reduction in Malat1 mRNA levels, indicating efficient pharmacological activity (Fig. 6j). To determine the cell-type specificity of this effect, we isolated endothelial and non-endothelial brain fractions using CD31-based magnetic bead separation. Quantitative analysis revealed that the Malat1–eExACT conjugate predominantly reduced Malat1 expression in endothelial cells, with minimal impact on non-endothelial populations (Fig. 6j). In contrast, administration of unconjugated Malat1 ASO led to mRNA reduction in both endothelial and non-endothelial compartments (Fig. 6j). These findings demonstrate that eExACT enables selective delivery of ASOs to brain endothelial cells *in vivo*, offering a promising strategy for targeted gene modulation while minimizing off-target effects of ASOs.

To probe the uptake mechanism underlying eExACT-based delivery, we next examined internalization of unconjugated eExACT in HEK293 cells expressing SLCO1A2, which showed efficient transporter-dependent uptake (Supplementary Figs. 14a and 14b). Given the relatively high molecular weight and hydrophilicity of eExACT and its conjugates, we hypothesized that uptake might involve endocytosis. Consistent with this, pharmacological inhibition with Pitstop 2 (clathrin-mediated endocytosis inhibitor) or Dynasore (dynamin inhibitor) caused a marked reduction in intracellular accumulation of eExACT (Supplementary Figs. 14a and 14b). Similarly, uptake of ASO-eExACT-3 conjugate was reduced by Dyansore treatment (Supplementary Figs. 14c–14d). These findings indicate that endocytosis is a key component of SLCO1A2-dependent uptake of the eExACT scaffold and likely contributes to the internalization of larger eExACT conjugates.

### Cell-Specific Pharmacological Access via Gene Therapy–Mediated Ectopic Transporter Expression

To explore the broader applicability of our platform, we investigated whether eExACT uptake could be enabled in cells that do not naturally express the cognate solute carrier transporter. We employed AAV-mediated gene delivery in live mice (Figs. 7a–7c) and in utero electroporation (Supplementary Figs. 15a–15c) to ectopically express SLCO1A2 in cortical neurons, which do not normally express this transporter. Uptake of eExACT and the colchicine–eExACT conjugate was then assessed in transfected neurons using intravital two-photon imaging. Following topical cortical administration of the compounds, we observed robust and selective uptake in neuronal cell bodies (Figs. 7b– 7c) and dendrites (Supplementary Figs. 15b–15c). Neurons expressing SLCO1A2-GFP showed strong, cell-restricted accumulation of the conjugate, whereas neighboring GFP-only neurons displayed little to no fluorescence despite being exposed to the same extracellular concentration. This sharp contrast between transporter-positive and control neurons indicates that ectopic expression of SLCO1A2 is sufficient to gate neuronal entry of eExACT and eExACT-conjugated molecules *in vivo*. Interestingly, ectopic neuronal uptake was accompanied by a marked reduction in endothelial labeling, likely reflecting redistribution of the compounds into the expanded neuronal compartment (Fig. 7c and Supplementary Figs. 15b and 15c). Systemic administration of eExACT led to progressive accumulation of fluorescence within the brain interstitium over time (Supplementary Fig. 17b–17c), indicating that the compound can cross the blood–brain barrier. In separate experiments, systemic eExACT dosing also resulted in detectable accumulation in neurons that ectopically express the transporter (Supplementary Fig. 17a), consistent with transporter-dependent uptake once the compound has entered the brain.

**Fig. 7:**
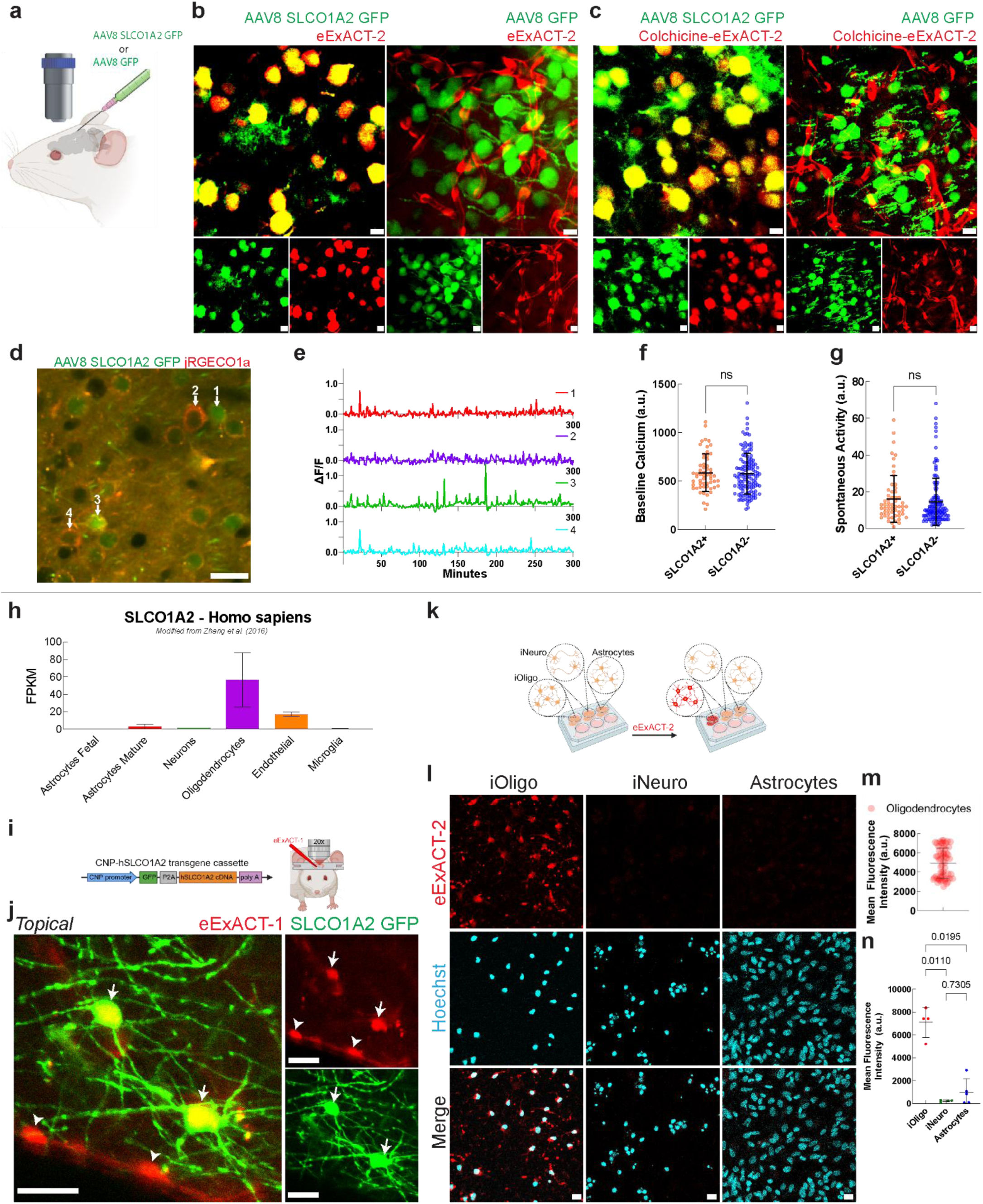
Human SLCO1A2 Enables In Vivo Cell Type-Specific eExACT Uptake in Neurons and Oligodendrocytes and in Human iPSC-Derived Oligodendrocytes. **a.** Schematic of AAV8-mediated intracortical delivery of either SLCO1A2-GFP or GFP control to drive cell type–specific expression of the human solute carrier SLCO1A2 in neurons. **b.** Two-photon *in vivo* images showing selective uptake of eExACT-2 (red, 50 µM) in neurons expressing SLCO1A2-GFP (green, left), with no uptake in control neurons expressing GFP only (right). Insets display separated fluorescence channels. Notice the loss of endothelial uptake with SLCO1A2 expression, likely due to competition for compound uptake. **c.** Two-photon images showing selective uptake of Colchicine-eExACT-2 (red, 50 µM) in SLCO1A2-GFP–positive neurons (left), but not in GFP-only controls (right). Endothelial labeling is similarly reduced in SLCO1A2-expressing brains. Scale bars for b and c, 10 μm. d. *In vivo* two-photon image showing co-expression of SLCO1A2-GFP (green) and the red calcium indicator iRGECO1a in cortical neurons. Numbered cells include both SLCO1A2-transfected (white arrow 1 and 3, GFP+) and non-transfected (white arrow 2 and 4, GFP−) neurons present in the same field of view. These neurons were imaged in parallel over time using time-lapse two-photon microscopy to assess calcium dynamics. *Scale bar, 20 μm.* e. Representative traces of spontaneous calcium activity (ΔF/F) from individual SLCO1A2-expressing (white arrow 1 and 3 from panel **d**, GFP+) and non-expressing (white arrow 2 and 4 from panel **d**, GFP−) cortical neurons imaged in the same field of view over a 300-minute time-lapse two-photon recording. Traces show dynamic calcium fluctuations, indicating that SLCO1A2 expression does not disrupt baseline neuronal activity. **f–g.** Quantification of baseline calcium levels (f) and spontaneous activity (g) reveals no significant difference (ns) between SLCO1A2+ and SLCO1A2− neurons, confirming that AAV-mediated SLCO1A2 expression does not alter calcium homeostasis. **h.** Bar graph depicting SLCO1A2 transcript expression levels (FPKM) across human CNS cell types, including astrocytes fetal, astrocytes mature, neurons, oligodendrocytes, endothelial, and microglia. Expression is predominantly enriched in oligodendrocytes relative to all other CNS cell types examined. Data were modified from Zhang et al.^44^. Data are presented as mean ± SD. **i.** Schematic of the CNP-hSLCO1A2 transgene cassette, consisting of a CNP promoter driving GFP-P2A-hSLCO1A2 cDNA followed by a poly-A signal, used to generate humanized transgenic mice expressing the SLCO1A2 transporter selectively in oligodendrocytes via microinjection and embryo transfer. Transgenic mice were subsequently used for intravital two-photon imaging following topical cortical application of eExACT-1**. j.** Representative intravital two-photon images of the cortex in CNP-hSLCO1A2 transgenic mice following topical application of eExACT-1 (red, 50 µM). SLCO1A2-expressing oligodendrocytes are visualized via GFP fluorescence (green, white arrows) and endothelial cell bodies (red, white arrowheads). Co-localization of eExACT-1 and GFP signal demonstrates selective uptake of eExACT-1 into SLCO1A2-expressing oligodendrocytes. Scale bars, 20 µm. **k.** Schematic of the in vitro uptake assay in which iPSC-derived oligodendrocytes (iOligo), iPSC-derived neurons (iNeuro), and human astrocytes were cultured separately and treated with eExACT-2 to assess cell-type-specific uptake. **l.** Representative fluorescence images of iOligo, iNeuro, and astrocytes following eExACT-2 treatment (red, top row) and Hoechst nuclear counterstain (cyan, middle row), with merged channels shown (bottom row). Robust eExACT-2 uptake is observed exclusively in iOligo, with minimal to no signal detected in iNeuro or astrocytes, demonstrating oligodendrocyte-selective internalization of eExACT-2 in vitro. Scale bars, 20 µm. **m.** Quantification of mean fluorescence intensity of eExACT-1 in GFP-positive oligodendrocytes of CNP-hSLCO1A2 transgenic mice (n=3) following topical cortical application of eExACT-1 (50 µM). Each dot represents an individual cell. Data are presented as mean ± SD (related to j). **n.** Quantification of mean fluorescence intensity of eExACT-2 across iOligo, astrocytes, and iNeuro. Each dot represents an individual replicate. Data are presented as mean ± SD. Statistical comparisons were performed using the two-stage linear step-up procedure of Benjamini, Krieger, and Yekutieli, with significance determined by the Kruskal–Wallis test; p-values indicated (iOligo vs. iNeuro p = 0.0110, iNeuro vs. Astrocytes p = 0.7305, iOligo vs. Astrocytes p = 0.0195, related to l).

We next asked whether AAV-mediated SLCO1A2 expression alters neuronal physiology. To address this, we performed *in vivo* calcium imaging in mice expressing SLCO1A2-GFP in a subset of neurons and compared their activity to that of neighboring untransfected neurons (Figs. 7d–7g). Neurons overexpressing SLCO1A2 maintained normal baseline calcium levels and spontaneous transient frequencies, indistinguishable from adjacent GFP-only cells. These data indicate that ectopic transporter expression, while sufficient to confer selective uptake of eExACT-based cargos, does not measurably disrupt neuronal function under these conditions.

Together, these results establish the feasibility of using gene therapy–mediated transporter expression to enable non-invasive, cell-specific delivery of pharmacological agents to otherwise inaccessible populations in the brain and beyond.

### SLCO1A2 expression in human oligodendrocytes supports eExACT-based delivery to clinically relevant cell type

The experiments above demonstrate that AAV-mediated expression of SLCO1A2 is sufficient to reprogram eExACT uptake to genetically defined neuronal populations in the mouse brain. To assess how this platform might translate to human CNS cell types and identify additional populations naturally poised for eExACT-based delivery, we next examined the endogenous distribution of SLCO1A2 in human brain tissue. Reanalysis of human RNA-sequencing datasets^44^ revealed that SLCO1A2 transcripts are strongly enriched in oligodendrocytes and brain endothelial cells (Fig. 7h). This expression pattern suggested that eExACT could provide a particularly efficient entry route into oligodendroglial and vascular compartments in the human brain.

Guided by this expression pattern, we generated oligodendrocyte-specific hSLCO1A2 transgenic mice using the CNP promoter to drive a cassette that co-expresses GFP and hSLCO1A2, with GFP reporting transporter-expressing cells that display characteristic oligodendrocyte morphology (Fig. 7i–7j). Intravital two-photon imaging of these mice following topical cortical application of eExACT-1 revealed robust co-localization of eEXACT-1 fluorescence with GFP-labeled oligodendrocyte cell bodies (Fig. 7j, white arrows), together with the expected endothelial uptake signal (Fig. 7j, white arrowheads). This pattern demonstrates strong uptake into SLCO1A2-expressing oligodendrocytes while preserving the normal endothelial labeling pattern. Quantification of mean fluorescence intensity across individual GFP-positive oligodendrocytes confirmed strong and consistent intracellular accumulation of eExACT-1 in these cells (Fig. 7m).

We next compared eExACT uptake across human CNS cell types using iPSC-derived oligodendrocytes (iOligo), iPSC-derived neurons (iNeuro), and primary human astrocytes (Fig. 7k). After exposure to eExACT-2, iOligo showed bright intracellular fluorescence, whereas iNeuro and astrocytes in the same fields exhibited little to no detectable signal (Fig. 7l and 7n), highlighting the potential of this platform to support concurrent oligodendroglial and vascular delivery in the human CNS.

Altogether, our data demonstrate that small-molecule delivery to defined cell types can be achieved by functionally coupling cargo uptake to specific solute carriers, such as SLCO1A2 in oligodendrocytes and vascular cells. By designing modular small molecules with affinity for selected transporters, and leveraging both endogenous and engineered transporter expression, this strategy enables selective intracellular access to genetically or natively specified cell populations in the CNS, retina, and potentially other organs. In doing so, it establishes a general framework that can, in principle, be adapted to diverse transporter–cargo pairs to reach cellular targets that are otherwise difficult to access with conventional delivery approaches.

## Discussion

The clinical effectiveness of many potent pharmacological agents is often limited by off-target toxicity, particularly in cell populations not directly involved in the disease process.^45, 46^ Despite this, few strategies currently exist to develop drugs capable of selectively targeting specific cell types while sparing others.^45^ Here we introduce Engineered Drugs with Transporter-Mediated Cell Type Specificity (ExACT), a strategy that enables the discovery and deployment of transporter-selective chemical motifs for conjugation to therapeutic compounds. In addition to small molecules, we show that ExACT can be extended to larger therapeutics, including antisense oligonucleotides, enabling selective intracellular delivery and functional activity in targeted cells. The marked reduction in systemic toxicity observed with these conjugates highlights the potential of ExACT to expand the therapeutic window and enable the repurposing of broadly acting agents,^47^ including those targeting widely expressed intracellular pathways.

Our discovery strategy was inspired by the observation that certain fluorescent compounds exhibited selective uptake into specific brain cell types *in vivo*^17–20^. We hypothesized that this selectivity was driven by chemical functional groups that engage membrane transporters enriched in distinct cell populations. To identify such functional groups in an unbiased manner, we synthesized a combinatorial library featuring diverse fluorophore scaffolds and varied chemical modifications, and performed large-scale two-photon imaging–based screening directly in the live mouse brain. This *in vivo* approach allowed us to uncover cell-type–selective uptake within the native brain microenvironment, without prior knowledge of the underlying transport mechanisms or the identity of the transporters involved. In future work, higher-throughput in vitro screens could be deployed once a relevant transporter has been identified or when focusing on human transporters, for example by using transporter-expressing cell lines, iPSC-derived cells, or organoids. In parallel, incorporating transporter-targeting motifs into drug-like small-molecule libraries could support high-throughput screening for both cell-type selectivity and pharmacological efficacy.

For our proof-of-principle conjugation experiments, we selected colchicine due to the extensive prior literature characterizing its essential functional groups and conjugation-tolerant sites^29–31^, allowing us to design bifunctional compounds that retain microtubule-targeting activity after conjugation. This enabled a direct demonstration that potent pharmacological effects can be preserved while adding a transporter-binding module. For less well-characterized or novel compounds, additional structure–activity analyses will likely be required to distinguish essential moieties from positions that can be modified. In such cases, the use of cleavable linkers could permit intracellular release of the native drug after transporter-mediated uptake, helping to maintain on-target activity while still conferring cell-type specificity.^48^

Regarding the types of therapeutic agents that can be delivered, our conjugation experiments encompassed both small molecules (up to approximately 1 kDa) and larger nucleic acid therapeutics, such as antisense oligonucleotides (ASOs), which showed robust, selective uptake and functional knockdown in brain endothelial cells *in vivo*. These results demonstrate that ExACT can support efficient intracellular delivery and target engagement for cargos beyond conventional small molecules. To probe the uptake mechanism for higher-molecular-weight cargos, we performed pharmacological inhibition experiments and found evidence for a key role of endocytosis. These data suggest that large cargos are internalized predominantly via transporter-coupled endocytosis rather than direct translocation across the membrane, supporting the adaptability of the ExACT concept for delivering other therapeutic agents such as peptides, proteolysis-targeting chimeras (PROTACs) and si/sgRNAs.^49–51^

The remarkable selectivity of the molecules we identified for distinct neural cell types, including endothelium, pericytes, astrocytes, neurons, and oligodendrocytes, suggests potential therapeutic applications across a broad spectrum of brain and retinal disorders. In future work, transporter-guided conjugation strategies could be developed to systematically exploit this broader transporter landscape. For example, dual-specificity ExACT conjugates could be designed to first cross the blood–brain barrier via endothelial uptake and then engage a second transporter on target cells such as astrocytes, neurons, or oligodendrocytes, enabling multi-step targeting within the CNS. Such approaches could greatly expand the therapeutic reach of ExACT by enabling selective delivery not only across tissue barriers but also into specific cellular compartments of the brain and retina.

Transporter expression patterns can vary substantially between species and across cell types, and these features need to be considered when designing ExACT-style conjugates for human use. At very high topical concentrations, some eExACT conjugates exhibited delayed, low-level uptake in astrocytes following their initial rapid accumulation in endothelial cells, indicating that astrocytes can serve as a minor sink under conditions of high exposure. Consistent with this, we also observed very low levels of uptake in HEK293 cells transfected with SLCO1C1, a transporter that is primarily expressed in human astrocytes, suggesting that related organic anion transporters can mediate weak eExACT entry outside the principal target population. Although the mechanisms underlying this secondary uptake remain to be fully defined, these findings highlight that even low-affinity interactions with additional transporters can become apparent at high doses and should be taken into account when optimizing concentration and dosing regimens.

The selectivity of our eExACT molecules for human SLCO1A2, together with our humanized mouse and iPSC data, highlights SLCO1A2 as a practical entry point for therapeutic targeting in both brain and retina. In transgenic mice expressing human SLCO1A2 in oligodendrocytes, eExACT compounds showed efficient uptake into myelinating cells while preserving endothelial labeling, establishing that a single human transporter can support concurrent access to glial and vascular compartments *in vivo*. Complementary experiments in human iPSC-derived oligodendrocytes demonstrated robust eExACT uptake compared to neurons and astrocytes, indicating that human oligodendrocytes are natively poised to internalize SLCO1A2-guided cargos. In the retina, SLCO1A2 expression in retinal pigment epithelium and vasculature suggests that the same chemistry could be used to target these populations with high precision. Antisense oligonucleotides are increasingly being explored for retinal and CNS indications but often suffer from narrow therapeutic windows due to broad tissue distribution and dose-limiting toxicity. By enabling selective delivery of ASOs to SLCO1A2-expressing cell types, alongside cell-type-specific delivery of small-molecule therapeutics, SLCO1A2-guided ExACT conjugates could help widen the therapeutic window and support interventions that address vascular, myelin, and RPE pathology across a range of CNS and retinal disorders.

Our proof-of-concept experiments demonstrated that neurons, which do not naturally express Slco1a4 or SLCO1A2, became highly permeable to eExACT drug conjugates when these transporters were ectopically expressed. This finding illustrates how gene therapy can confer selective pharmacological access to otherwise inaccessible cell types by introducing defined membrane transporters. In future work, our screening strategies could be extended to identify small molecules with affinity for transporters derived from evolutionarily distant species. By combining such orthogonal transporters with matching ExACT chemistries, gene therapy could be used to install artificial uptake gateways in selected cells, enabling precise delivery of designer pharmacological agents with minimal off-target uptake in non-transduced mammalian tissues.

Finally, the discovery of small molecules with intrinsic cell-type specificity opens opportunities for developing neuroimaging probes with enhanced target selectivity. In particular, fluorinated or iodinated derivatives (Fig. S8a–S8c) could serve as radiotracers for positron emission tomography (PET), enabling *in vivo* imaging of transporter activity.^52^ This strategy could yield PET probes that selectively report on SLCO1A2 or other transporter function in specific brain cell types. Notably, SLCO1A2 variants have been associated with increased risk for progressive supranuclear palsy, potentially via effects on oligodendrocyte health,^53–55^ and elevated SLCO1A2 expression has been observed in the brains of individuals with Alzheimer’s disease.^56^ The ExACT platform may therefore provide a route to transporter-specific radiotracers that serve as biomarkers for disease diagnosis, patient stratification, and monitoring of target engagement over time.

## Methods

### Mice

All animal procedures were approved by the Yale University Institutional Animal Care and Use Committee (IACUC). Male and female mice aged 1–3 months were used for all experiments. Wild-type C57BL/6J (JAX #000664) mice or Tie2-GFP transgenic mice (JAX #003658) were used for compound uptake studies. In a subset of experiments, Oatp1a/1b cluster knockout mice (Taconic #10707)^57^ were used. *In vivo* calcium imaging was performed on C57BL/6J mice, and CD-1 (Charles River #022) mice were used for antisense oligonucleotide experiments.

### Synthesis of Combinatorial Fluorescent Chemical Library

All starting materials were non-fluorescent and commercially available. To enhance chemical diversity and functionality, select starting materials were first converted into different salt forms. Reactions were typically carried out by dissolving the reactants (10 mM) and catalysts (10 µM) in 0.5 mL of dimethyl sulfoxide (DMSO) in 1-dram clear glass vials, followed by vortexing for 1 minute. To increase throughput, reactions (5 vials per round) were subjected to microwave irradiation (800 W, 3 minutes), yielding products with 20–95% efficiency. During the reaction, most volatile components evaporated, leaving DMSO as the residual solvent. Crude products were used without further purification. For *in vivo* screening, ten unpurified compounds were pooled in equal proportions to form each batch, which was diluted in PBS (1:15 v/v), vortexed for 2 minutes, centrifuged, and the supernatant collected for injection.

### Viral Production

An adeno-associated virus (AAV8) encoding SLCO1A2 was custom-produced by VectorBuilder. Briefly, the human SLCO1A2 coding sequence was cloned downstream of the CAG promoter in a pAAV8 expression vector, followed by a P2A self-cleaving peptide and EGFP to enable bicistronic expression. The construct (pAAV-CAG>hSLCO1A2:P2A:EGFP) was transfected into HEK293 cells for viral packaging. The resulting AAV8 particles were harvested and ultra-purified to a final titer exceeding 10^13^ genome copies per milliliter (GC/mL).

### AAV Brain Injections

AAV was delivered into the mouse subarachnoid space using a previously described protoco^58^. Mice were anesthetized with ketamine–xylazine, and a ∼1 mm diameter cranial window was made at coordinates 6 mm posterior and 3 mm lateral to bregma using a high-speed drill. A 30-gauge needle was then used to gently lift the thinned skull, exposing the dura. AAV stock was diluted 1:20 in sterile phosphate-buffered saline (PBS), adjusted to a final titter of 10^12^ genome copies/mL, and kept on ice until use. The viral solution was loaded into a Tygon tube connected to a polypropylene tip (outer diameter 70 µm), which was attached to a Hamilton syringe via a programmable syringe pump. The tip was inserted into the subarachnoid space and secured with cyanoacrylate glue. A total volume of 10 µL was injected at 0.2 µL/min. After injection, the tip was withdrawn, and the scalp incision was sutured. Mice recovered on a heating pad, and stable fluorescent expression was confirmed by imaging three weeks post-injection.

### In Utero Electroporation

In utero electroporation was performed at embryonic day 17 (E17) to selectively transfect developing layer II cortical neurons, following previously described procedures.^59^ Pregnant mice were anesthetized with ketamine (100 mg/kg) and xylazine (10 mg/kg), and the abdominal area was shaved, sterilized, and incised along the midline to expose the uterine horns. A pulled glass micropipette was used to inject ∼0.5 µL of plasmid DNA (1 µg/µL) into the lateral ventricle of each embryo using a Picospritzer II (General Valve). Electroporation was then conducted with tweezer-style electrodes (BTX Harvard Apparatus), delivering 4 pulses at 50 V with a duration of 50 ms and 1-second intervals. After the procedure, the embryos were gently returned to the abdominal cavity, and both muscle and skin layers were sutured.

### Cranial Window Preparation and In Vivo Two-Photon Imaging

One-month-old wild-type mice were anesthetized with ketamine (100 mg/kg) and xylazine (10 mg/kg) and placed on a 37 °C heating pad. Buprenorphine (0.1 mg/kg) and carprofen (5 mg/kg) were administered subcutaneously. The scalp was shaved and disinfected with povidone-iodine and ethanol, and ophthalmic ointment was applied to protect the eyes. A small region of scalp was removed to expose the skull. A circular craniotomy was performed by thinning and lifting the skull without damaging the underlying brain. The dura was gently removed using fine forceps, and a 4 mm cover glass was placed directly onto the brain and sealed to the skull. Fluorescent dyes or drug conjugates were administered either intravenously or topically for 20 minutes prior to window placement. A custom head bar was affixed to the skull using adhesive (for acute imaging) or dental cement (for chronic imaging). Imaging was performed using a two-photon microscope (Prairie Technologies) equipped with a mode-locked MaiTai tunable laser (Spectra Physics) and a 20x water-immersion objective (Zeiss, 1.0 NA). Imaging depths extended up to 300 µm below the pial surface, using excitation wavelengths between 750 and 950 nm, as previously described.^60^ For chronic imaging, mice were allowed to recover on a heated pad and received postoperative analgesia with buprenorphine and carprofen for 3 days.

### Combinatorial Fluorophore Batch Screening

Batches of 10 fluorescent compounds were topically applied to the cortical surface via the craniotomy immediately before placement of the cover glass. No wash steps were performed, allowing for continuous imaging of the exposed cortex for up to 3 hours using two-photon microscopy. Imaging data were analyzed to identify patterns of cell-type–specific labeling. Most batches displayed diffuse fluorescence in interstitial spaces or nonspecific cellular labeling. Batches that showed any indication of selective labeling were prioritized for further analysis. These “positive” batches were sequentially subdivided and imaged through three additional rounds of screening to isolate the compound responsible for the selective uptake pattern. Because the initial chemical mixtures were not purified, observed labeling could result from fluorescent byproducts or minor reaction components. Therefore, candidate compounds were resynthesized at larger scale, purified, and structurally characterized. The purified compounds were then re-evaluated *in vivo* to confirm reproducible, cell-type–specific labeling.

### *In Vitro* and *In Vivo* Compound Uptake Experiments

For *in vitro* assays, HEK293 cells were cultured in 24-well plates to approximately 60% confluency. Cells were transiently transfected with 250 ng of a candidate transporter plasmid using 1 µL of JetPRIME transfection reagent per well. After a 48-hour incubation period, cells were washed with PBS and incubated with novel fluorescent probes diluted in transport buffer at various concentrations. Following a 30-minute incubation at 37 °C, uptake was halted by washing the cells with PBS, followed by fixation with 4% paraformaldehyde. Transporter expression was detected by immunostaining against a FLAG epitope using an anti-FLAG Alexa Fluor 488 antibody. Cell nuclei were counterstained with Hoechst 33342. Samples were mounted and imaged using a Leica SP8 confocal microscope to assess uptake and transporter localization. For *in vivo* biodistribution studies, eExACT compounds (0.1 mM, 70 µL) were administered via tail vein injection in Tie2-GFP transgenic mice, which label endothelial cells. After 2.5 hours, mice were transcardially perfused, and organs including brain, spleen, heart, skeletal muscle, liver, and kidney were harvested, cryosectioned, and imaged by confocal microscopy. Endothelial-specific uptake and compound distribution in parenchymal tissues were evaluated semi-quantitatively across organs.

### Preparation of Acute Brain Slices

Acute brain slices were prepared from mice between postnatal days 20 and 30 (P20-P30). Slices were cut in the sagittal plane with a thickness of 200 µm at 1.2 mm lateral from the midline. They were incubated in artificial cerebrospinal fluid (ACSF) containing eExACT-2 (10 µM). The ACSF composition was as follows (in mM): 120 NaCl, 3.1 KCl, 1.1 CaCl_2_, 1.2 MgCl_2_, 1.25 MgSO_4_, 26 NaHCO_3_, 0.5 L-glutamine, 0.1 ascorbic acid, 0.1 Na-pyruvate, and 20 glucose. The ACSF was saturated with 95% O_2_ and 5% CO_2_ and maintained at 35°C. After a 10-minute incubation period, the slices were transferred to fresh ACSF. Two-photon imaging was then performed on different brain regions of the slices immediately following the transfer. This method ensures that the brain slices remain viable and suitable for imaging studies.

### Assessment of Pharmacological Effects In Vitro

NIH3T3 cells, which naturally take up eExACT compounds, and are commonly used for studies of microtubule polymerization, were employed to assess the pharmacological activity of the Colchicine–eExACT conjugate. Cells were treated for 12 hours with varying concentrations of Colchicine–eExACT, unconjugated Colchicine, or vehicle control. Confocal imaging of Hoechst-stained nuclei was used to quantify the proportion of multinucleated cells, a hallmark of Colchicine-induced mitotic disruption^61^. Additionally, immunofluorescence labeling with anti–α-tubulin antibodies was performed to evaluate changes in microtubule organization as a functional readout of drug activity.

### Cell-Free Tubulin Polymerization Assay

To evaluate the effects of eExACT compounds on microtubule dynamics, we performed a one-step fluorescence-based tubulin polymerization assay (Cytoskeleton, Inc., #BK011P) as previously described^32^. The assay was carried out at 39 °C in a pre-warmed 96-well plate with continuous orbital shaking to promote polymerization. Fluorescence intensity (λ_ex_ = 350 nm and λ_em_ = 435 nm) was recorded using an Infinite 200Pro plate reader (Tecan, i-control software) for up to 30 minutes following the addition of Colchicine–eExACT, unconjugated Colchicine, vehicle, or control eExACT compounds (3 µM).

### Assessment of In Vivo Compound Toxicity

To assess toxicity, Colchicine–eExACT, unconjugated eExACT, and Colchicine were administered via both systemic intraperitoneal injection or localized intradermal injection at varying concentrations. For systemic toxicity evaluation, we monitored weight daily in P20 mice during their active growth phase. For local toxicity, we used a hair regrowth assay: fur was shaved to stimulate follicular stem cell proliferation, and compounds were injected intradermally into the lower right back quadrant, with the contralateral left quadrant serving as a control. Hair regrowth was documented over 12 days using serial photography. Pixel intensity across the injection sites was quantified using the Plot Profile plug-in in NIH ImageJ. Local toxicity was expressed as the percent difference in pixel intensity between injected and non-injected regions using the formula:

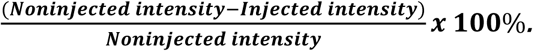

### Tissue Processing and Confocal Imaging

Brains were post-fixed overnight at 4 °C in 4% paraformaldehyde and sectioned at 50 µm using a vibratome (Leica VT1000). A Leica SP8 confocal microscope was used for high-resolution imaging, employing appropriate laser excitation lines and acousto-optical beam splitter settings to optimize fluorophore excitation and emission detection. The microscope was equipped with either a 20x water immersion objective (1.0 NA, Leica) or a 63x oil immersion objective (1.40 NA, Leica).

### Antisense Oligonucleotide (ASO) Synthesis and Purification

All antisense oligonucleotides (ASOs) were synthesized by BioSynthesis, Inc. using standard solid-phase phosphoramidite chemistry on controlled pore glass (CPG) supports. Synthesis was performed in the 3’ to 5’ direction using a DNA/RNA synthesizer under anhydrous conditions. Oligonucleotides were assembled with standard protecting groups and coupling cycles to ensure high coupling efficiency. Following synthesis, the oligonucleotides were deprotected and cleaved from the solid support using concentrated ammonium hydroxide or other appropriate deprotection reagents, depending on the sequence and base modifications. After deprotection, crude oligonucleotides were desalted and purified by reverse-phase high-performance liquid chromatography (RP-HPLC) or ion-exchange chromatography, as appropriate. For fluorophore labeling, oligonucleotides containing a terminal amine modification were reacted post-synthetically with NHS ester-activated fluorophores (e.g., Cy5) or sulfonyl chloride (eExACT-3). Labeling reactions were carried out in anhydrous dimethylformamide (DMF) with triethylamine to facilitate coupling. Labeled oligonucleotides were subsequently purified by HPLC to remove unreacted dye and other byproducts. All oligonucleotide products were analyzed by mass spectrometry (MALDI-TOF or ESI-MS) and analytical HPLC to confirm the correct molecular weight and assess purity.

### In Vivo Evaluation of Antisense Oligonucleotide (ASO) Activity

To assess cell-type-specific pharmacodynamic effects, 2.5 nmol of Malat1-eExACT-3 ASO, Malat1 ASO, and Control ASO were each administered via subarachnoid injection as previously described.^62^ Three days later, mice were perfused with ice-cold PBS. The diencephalon was removed, and the remaining brain tissue was processed for analysis. Cell-enriched fractions, including total cells, CD31⁺ endothelial cells, and CD31⁻ non-endothelial cells, were isolated using the Adult Brain Dissociation Kit (Miltenyi) followed by CD31 magnetic bead separation, as previously described.^63^ One-tenth of the dissociated cell suspension was reserved as the total fraction and mixed immediately with Trizol (Invitrogen). The remainder was processed to isolate CD31⁺ and CD31⁻ fractions, to which 100 μL of Trizol was added. RNA extraction was performed per the manufacturer’s instructions, and gene expression was quantified using the Luna Universal One-Step RT-qPCR Kit (NEB).

### RT-PCR Primer Sequences

The following primers were used for RT-PCR analysis. For the 18s ribosomal RNA gene: forward primer (18s-f) – GTAACCCGTTGAACCCCATT, and reverse primer (18s-r) – CCATCCAATCGGTAGTAGCG. For the Malat1 gene: forward primer (Malat1-f) – AGCAGGCATTTGTGAGAGGA, and reverse primer (Malat1-r) – ATGTTGCCGACCTCAAGGAA.

### Generation of CNP-EGFP-P2A-hSLCO1A2 transgenic mice

To achieve oligodendrocyte-selective expression of human SLCO1A2, we cloned an EGFP-P2A-hSLCO1A2 cassette downstream of the 2′,3′-cyclic nucleotide 3′-phosphodiesterase (CNP) promoter in the pCNP expression vector^64^ (kindly provided by Q. Richard Lu, Cincinnati Children’s Hospital Medical Center), which contains a human β-globin polyadenylation signal. The linearized CNP-EGFP-P2A-hSLCO1A2 fragment was excised by NotI digestion, gel-purified, and microinjected into pronuclei of fertilized (C57BL/6J × SJL/J) F2 zygotes, which were then implanted into pseudopregnant females using standard procedures. Genomic DNA from tail biopsies was screened by PCR using primers spanning the CNP promoter and EGFP coding sequence (forward: 5′-AGGGTGCTTTAAGTTAGGCTTGGG-3′; reverse: 5′-ACACGCTGAACTTGTGGCCGTTTA-3′). PCR-positive founders were crossed to (C57BL/6J × SJL/J) F2 wild-type mice to confirm germline transmission, and hemizygous offspring were intercrossed to establish stable CNP-EGFP-P2A-hSLCO1A2 lines. EGFP expression was assessed by fluorescence imaging in acute and fixed brain preparations, confirming selective labeling of CNP-positive oligodendrocytes with no detectable expression in neurons, astrocytes, or microglia. Transgenic mice were viable, fertile, and displayed no overt anatomical or behavioral abnormalities under standard housing conditions. All animal procedures were approved by the Yale Institutional Animal Care and Use Committee and complied with institutional and national guidelines.

### Human Cell Culture

#### Human iPSC-derived oligodendrocyte-like cells

Human iPSC-derived oligodendrocyte-like cells (iOLCs; bit.bio, cat. no. io1028S) were thawed and seeded at 1.5 × 10⁴ cells per well in black-walled, glass-bottom 96-well plates (Cellvis) pre-coated with 1× Geltrex (Thermo Fisher Scientific). Cells were initially maintained for 24 h in oligodendrocyte differentiation medium (OM1), consisting of DMEM/F-12 supplemented with 1× B-27, 1× N-2, 1× GlutaMAX, 1× MEM non-essential amino acids (NEAA), 13.6 µM β-mercaptoethanol, 7 µM insulin, 60 ng mL⁻¹ triiodothyronine (T3), 100 ng mL⁻¹ biotin, 100 µM dibutyryl cAMP, 20 ng mL⁻¹ PDGF-AA, 5 ng mL⁻¹ basic FGF, 100 nM retinoic acid, 1 µM purmorphamine, 1 µg mL⁻¹ doxycycline, 5 µM ROCK inhibitor (Y-27632), and 1 µg mL⁻¹ laminin. Medium was then replaced with oligodendrocyte maturation medium (OM2), consisting of DMEM/F-12 supplemented with 1× B-27, 1× N-2, 1× GlutaMAX, 1× MEM NEAA, 13.6 µM β-mercaptoethanol, 7 µM insulin, 60 ng mL⁻¹ T3, 100 ng mL⁻¹ biotin, 100 µM dibutyryl cAMP, 5 ng mL⁻¹ NT-3, 10 ng mL⁻¹ IGF-1, 1 µg mL⁻¹ doxycycline, 5 µM ROCK inhibitor, and 1 µg mL⁻¹ laminin. Half-medium changes were performed every other day, and cells were used for live-cell imaging and immunocytochemistry between days in vitro (DIV) 7 and DIV 10.

#### Human primary astrocytes

Human primary astrocytes (ScienCell, cat. no. 1800) were thawed and maintained in complete Astrocyte Medium (ScienCell, cat. no. 1801) containing 1× fetal bovine serum (FBS), 1× Astrocyte Growth Supplement, and 1× penicillin–streptomycin, with 5 µM ROCK inhibitor added during all passaging and seeding steps. Cells at passage 2 or 3 were seeded at 1.5 × 10⁴ cells per well onto Geltrex-coated black-walled, glass-bottom 96-well plates. Half-medium changes were performed every other day, and cells were used for live-cell imaging and immunocytochemistry between DIV 14 and DIV 30.

#### Human NGN2-induced neurons

Human induced neurons were generated by doxycycline-inducible overexpression of Neurogenin-2 (NGN2) in KOLF2.1J iPSCs and were obtained from the Yale iPSC NeuroCore. Briefly, iPSCs were engineered to carry a stably integrated, doxycycline-inducible human NGN2 cassette using a piggyBac (PB) transposon strategy. The human NGN2 coding sequence under tetracycline operator (TetO) control (Addgene, plasmid no. 172115) was cloned into a piggyBac donor vector (PB-TO-hNGN2; VectorBuilder). Stable genomic integration was achieved by nucleofection of iPSCs with PB-TO-hNGN2, which encodes the doxycycline-inducible NGN2 cassette and a puromycin resistance gene, together with a transient piggyBac transposase expression plasmid (pRP[Exp]-EGFP/Hygro-CAG>PBase; VectorBuilder) encoding PBase and an EGFP–Hygromycin marker. The transposase plasmid was transiently expressed to mediate donor transposition and was not designed for stable integration; any transient GFP signal observed immediately post-nucleofection was attributable to this plasmid and diminished as it was diluted over subsequent passages. Cells were selected with puromycin to enrich for clones with stable PB-TO-hNGN2 integration. For neuronal differentiation, Geltrex-coated black-walled, glass-bottom 96-well plates were seeded at 1.5 × 10⁴ cells per well in neuronal induction medium (NIM) consisting of DMEM/F-12 supplemented with 1× GlutaMAX, 1× sodium pyruvate, 1× N-2, 1× B-27 minus vitamin A, 1 µg mL⁻¹ doxycycline, and 5 µM ROCK inhibitor. NIM was replaced daily for 3 days. On day 3, cultures were transitioned to BrainPhys Neuron Medium (STEMCELL Technologies, cat. no. 05790) supplemented with 1% N-2, 2% B-27 minus vitamin A, 1 µg mL⁻¹ natural mouse laminin (Thermo Fisher Scientific, cat. no. 23017015), 20 ng mL⁻¹ BDNF (R&D Systems, cat. no. 248), 20 ng mL⁻¹ GDNF (R&D Systems, cat. no. 212), 250 µg mL⁻¹ dibutyryl cAMP (Sigma-Aldrich, cat. no. D0627), and 200 µM L-ascorbic acid (Sigma-Aldrich, cat. no. A4403); half-medium changes were performed every other day for 10 days. Cells were subsequently maintained in Neuronal Maintenance Medium (NMM) consisting of 1× BrainPhys basal medium (STEMCELL Technologies), 1× B-27 with vitamin A (Thermo Fisher Scientific), 1× N-2 (Thermo Fisher Scientific), 5 µg mL⁻¹ cholesterol (Sigma-Aldrich), 1 mM creatine (Sigma-Aldrich), 10 nM β-estradiol, 200 nM ascorbic acid, 1 mM cAMP (Sigma-Aldrich), 20 ng mL⁻¹ BDNF (PeproTech), 20 ng mL⁻¹ GDNF (PeproTech), 1 µg mL⁻¹ laminin, 0.5 mM GlutaMAX (Thermo Fisher Scientific), 1 ng mL⁻¹ TGF-β1 (PeproTech), 1× antibiotic–antimycotic, and 50 U mL⁻¹ penicillin–streptomycin (Thermo Fisher Scientific). Half-medium changes were performed every other day, and neurons were used for experiments between DIV 20 and DIV 30.

### Live Human Cell Imaging of eExACT-2 Uptake

Live-cell imaging of eExACT-2 uptake was performed in human iPSC-derived oligodendrocyte-like cells, human primary astrocytes, and human NGN2-induced neurons at the Yale Neuroscience Imaging Core Facility using a Zeiss LSM 880 laser-scanning confocal microscope equipped with an environmental chamber maintained at 37°C and 5% CO₂. Prior to imaging, cells were washed once with phosphate-buffered saline (PBS) and incubated with Hoechst 33342 (1:2,000 dilution) for 10 min at 37°C. Cells were then washed twice with PBS and returned to their respective maintenance media. Baseline fluorescence images were acquired before eExACT-2 addition using defined laser power and detector gain settings, which were held constant throughout all subsequent acquisitions. A stock solution of eExACT-2 was prepared at 100 mM in dimethyl sulfoxide (DMSO) and diluted to a final working concentration of 100 µM in the appropriate cell culture medium immediately before use. Following 30 min of incubation at 37°C, cells were washed five times with PBS and imaged under the same acquisition parameters as baseline. For quantification of eExACT-2 uptake, Hoechst 33342 fluorescence was used to segment nuclei and define individual cell regions of interest (ROIs). Each nuclear ROI was radially expanded by a factor of 1.5 to approximate the whole-cell area. ROIs with areas below 7 pixels were excluded from analysis. The median integrated fluorescence density in the red channel was calculated for each ROI, background-corrected by subtracting the corresponding baseline fluorescence value, and normalized to the number of nuclei per field of view.

### Immunocytochemistry of Human Cells

Immunocytochemistry was performed on human iPSC-derived oligodendrocyte-like cells, human primary astrocytes, and human NGN2-induced neurons. To minimize mechanical disturbance while ensuring efficient solution exchange, all wash steps were performed using 75% volume exchanges and all antibody incubation steps using 90% volume exchanges. Cells were washed twice with ice-cold PBS and fixed with 4% paraformaldehyde (PFA) in PBS for 10 min at room temperature. Following fixation, cells were washed twice with PBS and permeabilized with 0.02% saponin in PBS for 15 min at room temperature. Cells were then washed twice with PBS and blocked with 10% normal goat serum (NGS) in PBS for 30 min at room temperature to reduce non-specific antibody binding. Primary antibodies (see Supplementary Information) were diluted in blocking buffer and incubated overnight at 4°C. Following primary antibody incubation, cells were washed five times with PBS and incubated with fluorophore-conjugated secondary antibodies (see Supplementary Information) for 1 h at room temperature, protected from light.

## Acknowledgements

This project was supported by National Institutes of Health grants R01NS111961 (JG), R01NS115544 (JG and RWG), and RF1NS128953 (JG and RWG); the Yale Center for Clinical Investigation (YCCI) CTSA Grant Number UL1TR001863 (RWG); and the Wu Tsai Institute Summer Undergraduate Program (S.P.). We thank the iPSC Core Facility led by Dr. Tanina Arab at the Yale Department of Neuroscience for providing stem cell-derived materials. This work is supported in part by the Yale Alzheimer’s Disease Research Center (ADRC, P30 #AG066508 awarded to Dr. Stephen Strittmatter and Dr. In-Hyun Park). Several diagrams in this manuscript were created using BioRender software.

## Author Contributions

J.G. and R.W.G. conceived the project. R.W.G. designed and performed the chemical synthesis of the combinatorial library and compound conjugations, with input from J.G. L.Z. designed and conducted experiments related to the identification and characterization of membrane transporters, with input from J.G. and R.W.G. J.Z. designed and implemented experiments assessing ASO target engagement *in vivo* and *in vitro*. J.Z. and E.S. implemented human iPSC experiments. H.K.T. and S.P. assisted with membrane transporter identification experiments, cell culture, intravital two-photon imaging, immunofluorescence confocal imaging, and studies of drug efficacy and toxicity. B.N. and J.C. contributed to intravital two-photon compound screening. L.T. performed intravital calcium and structural brain imaging, brain slice experiments, in utero electroporation, and subarachnoid viral injections, and provided microscopy technical support and quantitative image analysis. J.G. and R.W.G. wrote the manuscript. J.G. directed the overall project.

---

**Supplementary Figure 1.**
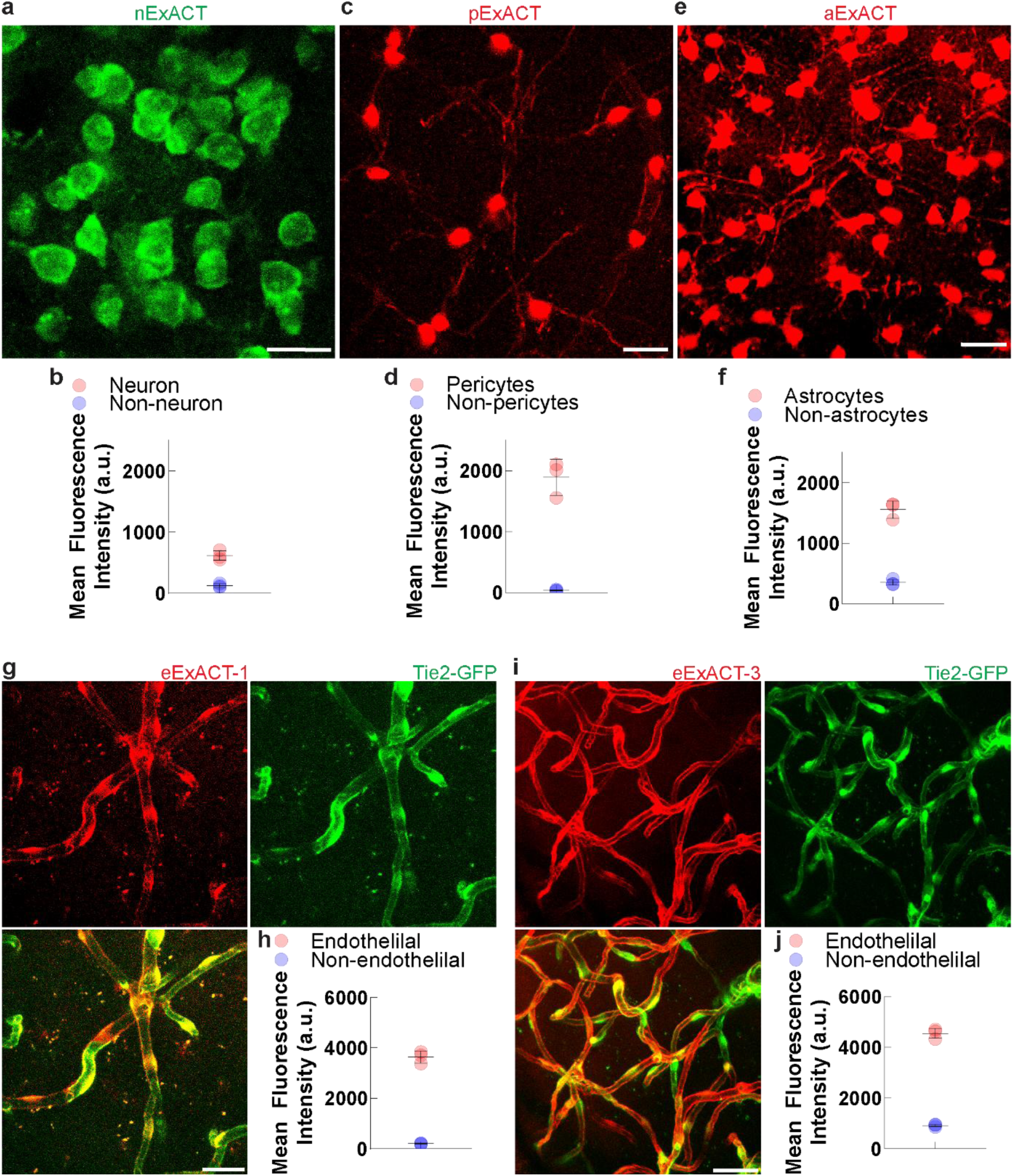
(related to Fig. 1). Cell-type–selective fluorophores demonstrate preferential labeling of neurons, pericytes, astrocytes, or endothelial cells. **a, c, e.** Representative two-photon *in vivo* images showing selective labeling by distinct fluorophores: nExACT (a) labels neurons (green), pExACT (c) labels pericytes (red), and aExACT (e) labels astrocytes (red). Scale bars, 20 μm. **b, d, f.** Quantification of mean fluorescence intensity comparing labeled versus non-labeled cell populations for nExACT in neurons (b), pExACT in pericytes (d), and aExACT in astrocytes (f). **g, i.** Intravital two-photon images of mouse cortex showing selective endothelial uptake of eExACT-1 (g) and eExACT-3 (i) (red), visualized in Tie2-GFP mice (green). Bottom panels show merged channels (yellow). Scale bars, 20 μm. **h, j.** Quantification of fluorescence intensity in endothelial versus non-endothelial structures 30 minutes after topical application of eExACT-1 (h) and eExACT-3 (j) confirms high cell-type specificity. All compounds were applied at 50 µM unless otherwise indicated.

**Supplementary Figure 2.**
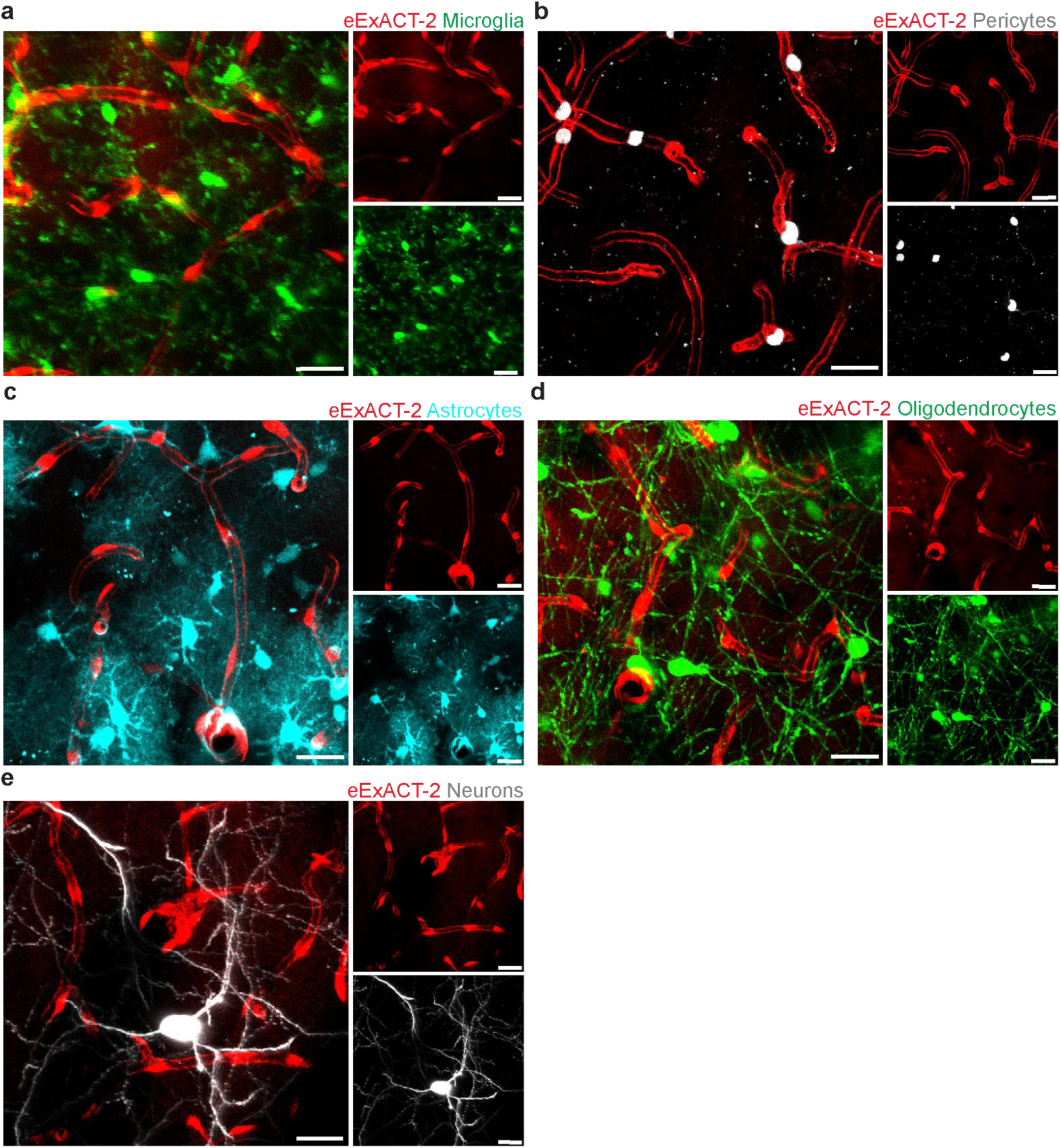
(Related to Fig. 1). Intravital two-photon imaging reveals selective endothelial uptake of eExACT-2 with no detectable uptake by other major brain cell types. **a-e.** Following topical application of eExACT-2 (red), cortical imaging in CX3CR1-GFP mice showed no colocalization with microglia (green, a). Pericytes labeled with Neurotrace (gray) similarly lacked eExACT-2 uptake (b). Astrocytes labeled via AAV-PHP.eB-GFAP-GFP (cyan) showed no overlap with eExACT-2 signal (c). Oligodendrocytes in PLP-eGFP mice (green) (d) and neurons in Thy1-GFP mice (gray) (e) also exhibited no colocalization with eExACT-2. Insets in right panels show individual fluorescence channels. Scale bars, 20 μm. All compounds were applied at 50 μM unless otherwise indicated.

**Supplementary Figure 3.**
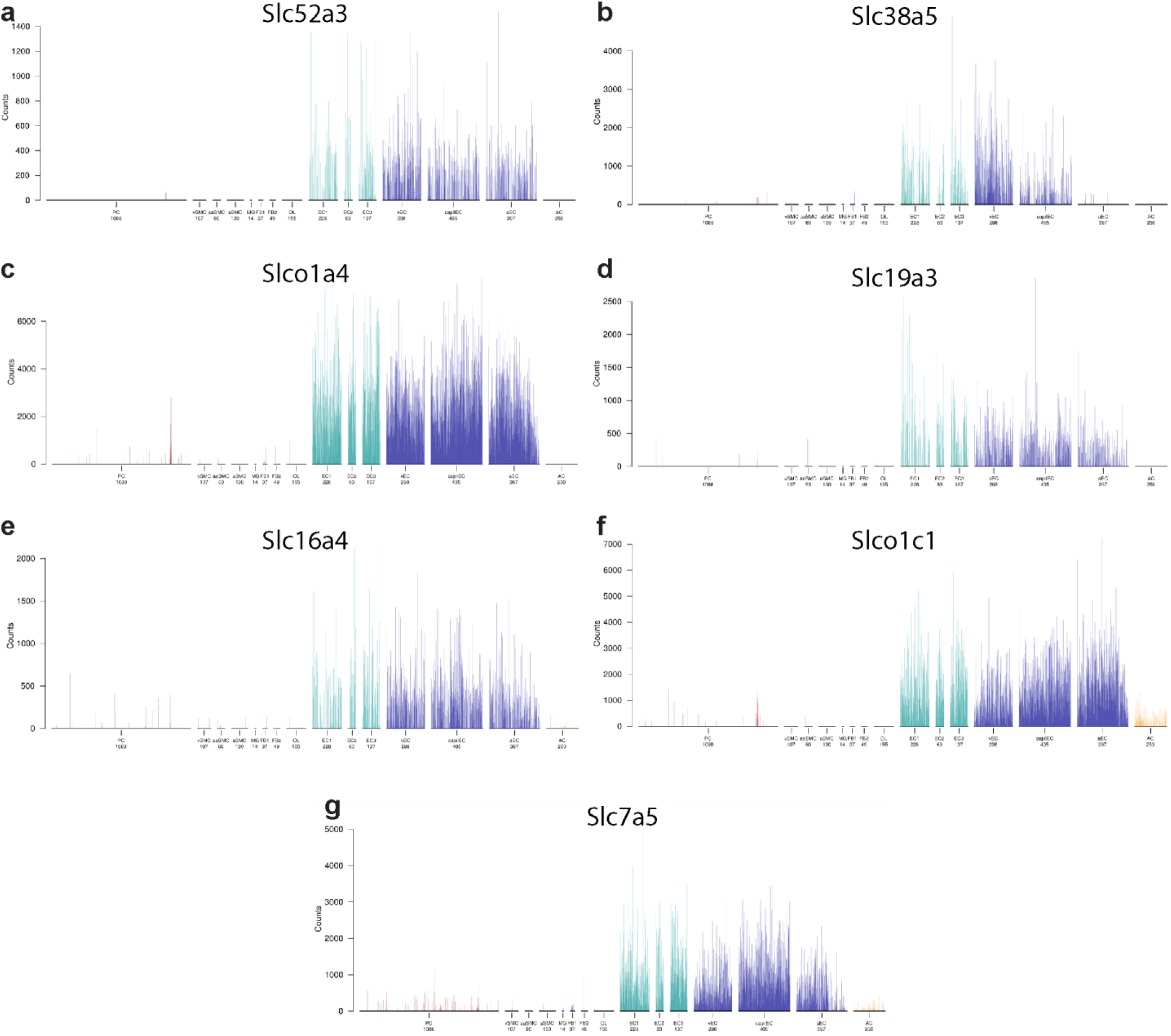
(Related to Fig. 2). Single-cell transcriptomic data showing solute carrier transporters with selective expression in brain endothelial cells. **a–g.** Expression patterns of selected solute carrier transporter genes in single-cell RNA-sequencing datasets from the mouse brain vasculature atlas. Genes shown include *Slc52a3* (a), *Slc38a5* (b), *Slco1a4* (c), *Slc19a3* (d), *Slc16a4* (e), *Slco1c1* (f), and *Slc7a5* (g). These genes display high expression in endothelial cell populations with minimal to no expression in other brain cell types. Data adapted from Vanlandewijck et al., *Nature* 2018^25^ .

**Supplementary Figure 4.**
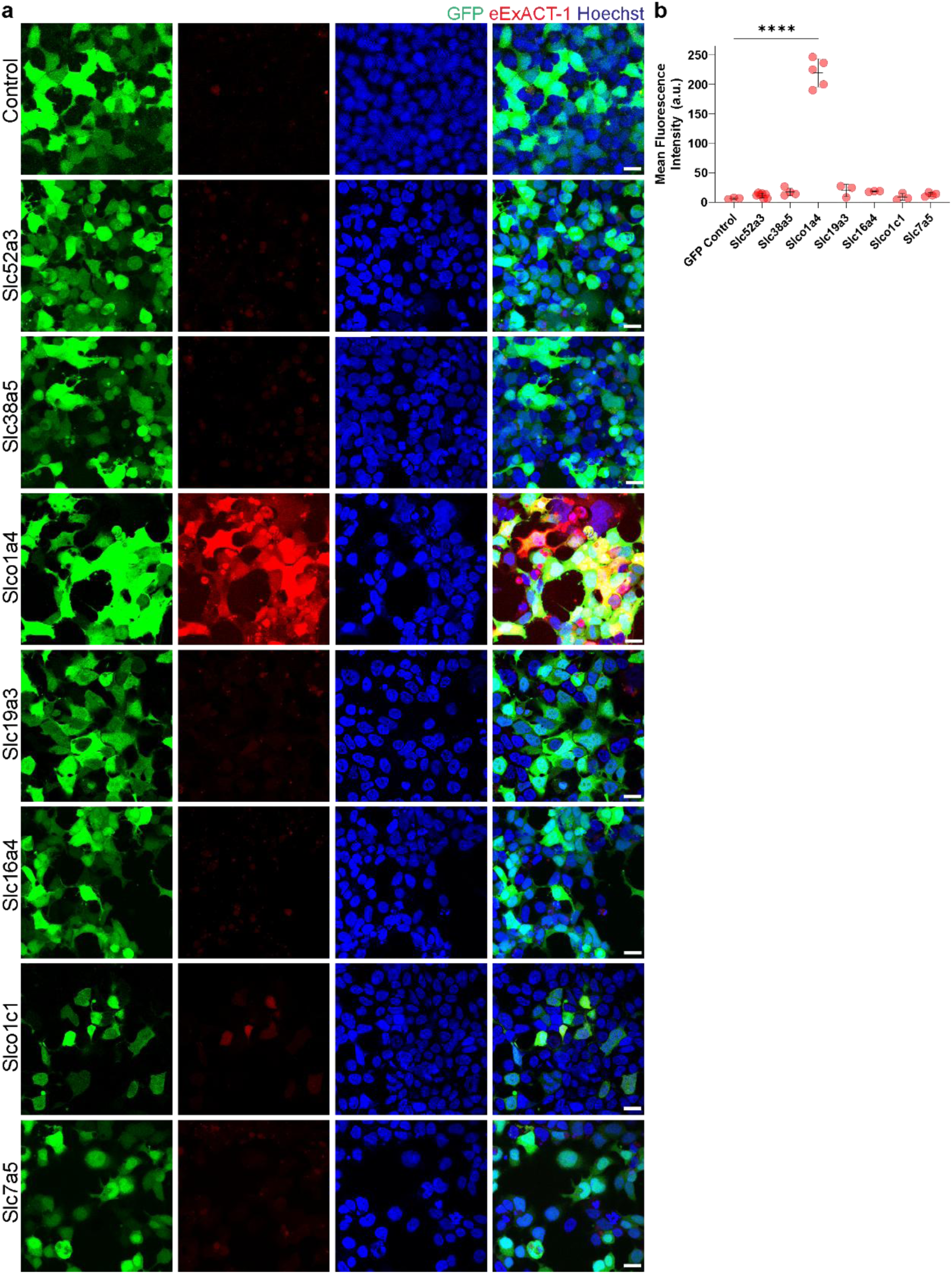
(Related to Fig. 2). Analysis of eExACT uptake upon transfection of various solute carrier transporters in HEK293 Cells. **a.** Evaluation of confocal images of HEK293 cells transfected with the solute carriers Slc7a5, Slco1c1, Slc16a4, Slc19a3, Slco1a4, Slc38a5, Slc52a3, or GFP control following exposure to eExACT-1 (red,10 μM) and nuclear stain (blue, Hoechst). Robust intracellular uptake is observed only in cells expressing the solute carrier Slco1a4, indicating its role as the eExACT-1 uptake transporter in the mouse brain. Scale bars, 20 μm. **b.** Quantification of mean fluorescence intensity (MFI) in HEK293 cells transfected with the solute carriers Slc7a5, Slco1c1, Slc16a4, Slc19a3, Slco1a4, Slc38a5, Slc52a3, or GFP control, after eExACT-1 (red, 10 μM) exposure (related to a). A statistical comparison was performed using two-tailed unpaired t-test where **** indicates P<0.001.

**Supplementary Figure 5.**
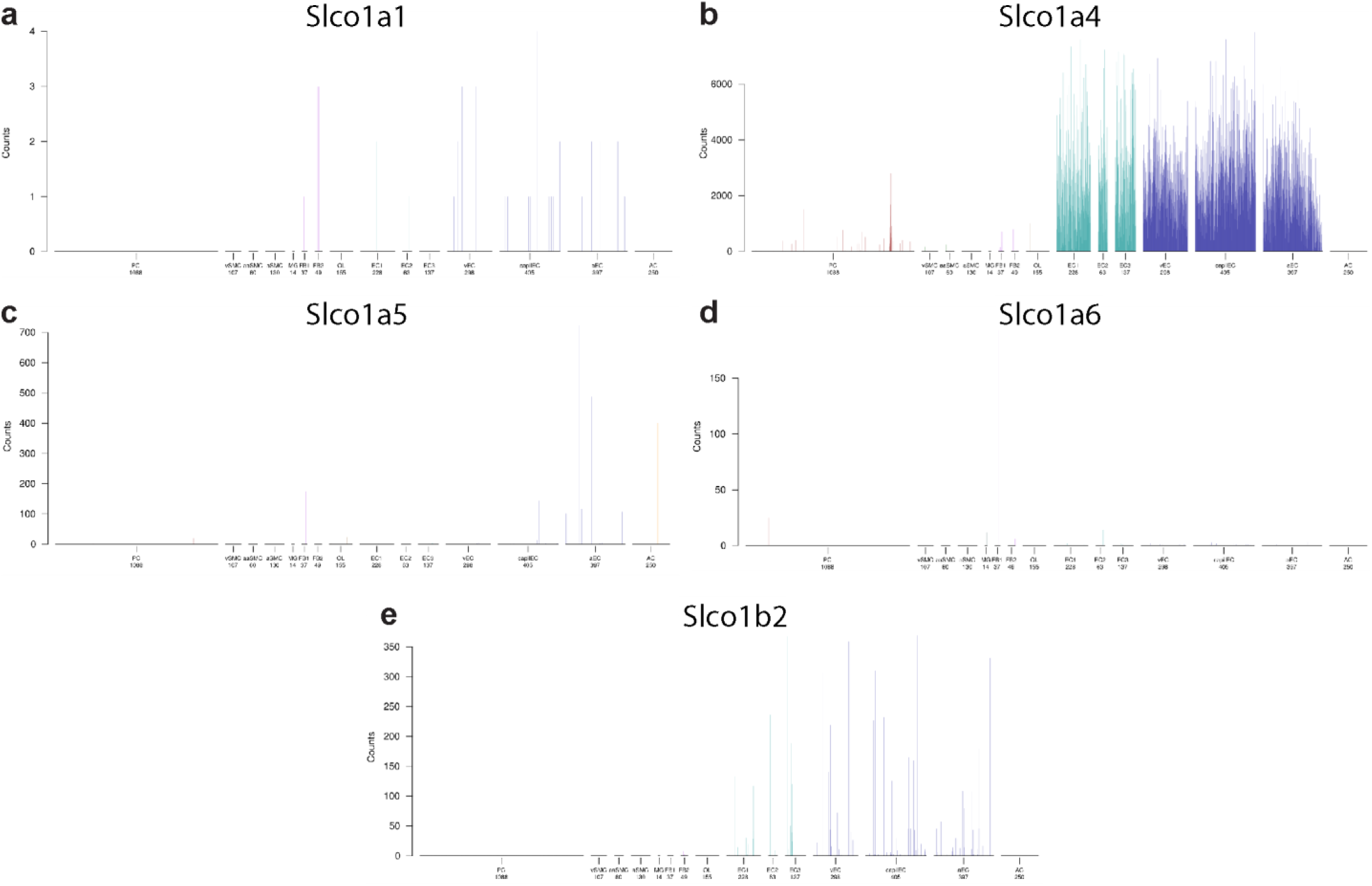
(Related to Fig. 2). Expression patterns of genes deleted in the Oatp1a/1b cluster knockout mouse, based on single-cell RNA sequencing of the mouse brain. **a–e.** Expression profiles of Slco genes deleted in the Oatp1a/1b cluster knockout mouse model: *Slco1a1* (a), *Slco1a4* (b), *Slco1a5* (c), *Slco1a6* (d), and *Slco1b2* (e). This mouse strain carries a deletion of five annotated *Slco1a* and *Slco1b* genes, as well as two predicted *Slco1a*-like genes. Among these, only *Slco1a4* (b) is detectably expressed in the brain and is selectively enriched in endothelial cells, supporting its role in brain-specific transport. Data from Vanlandewijck et al., *Nature* 2018^25^.

**Supplementary Figure 6.**
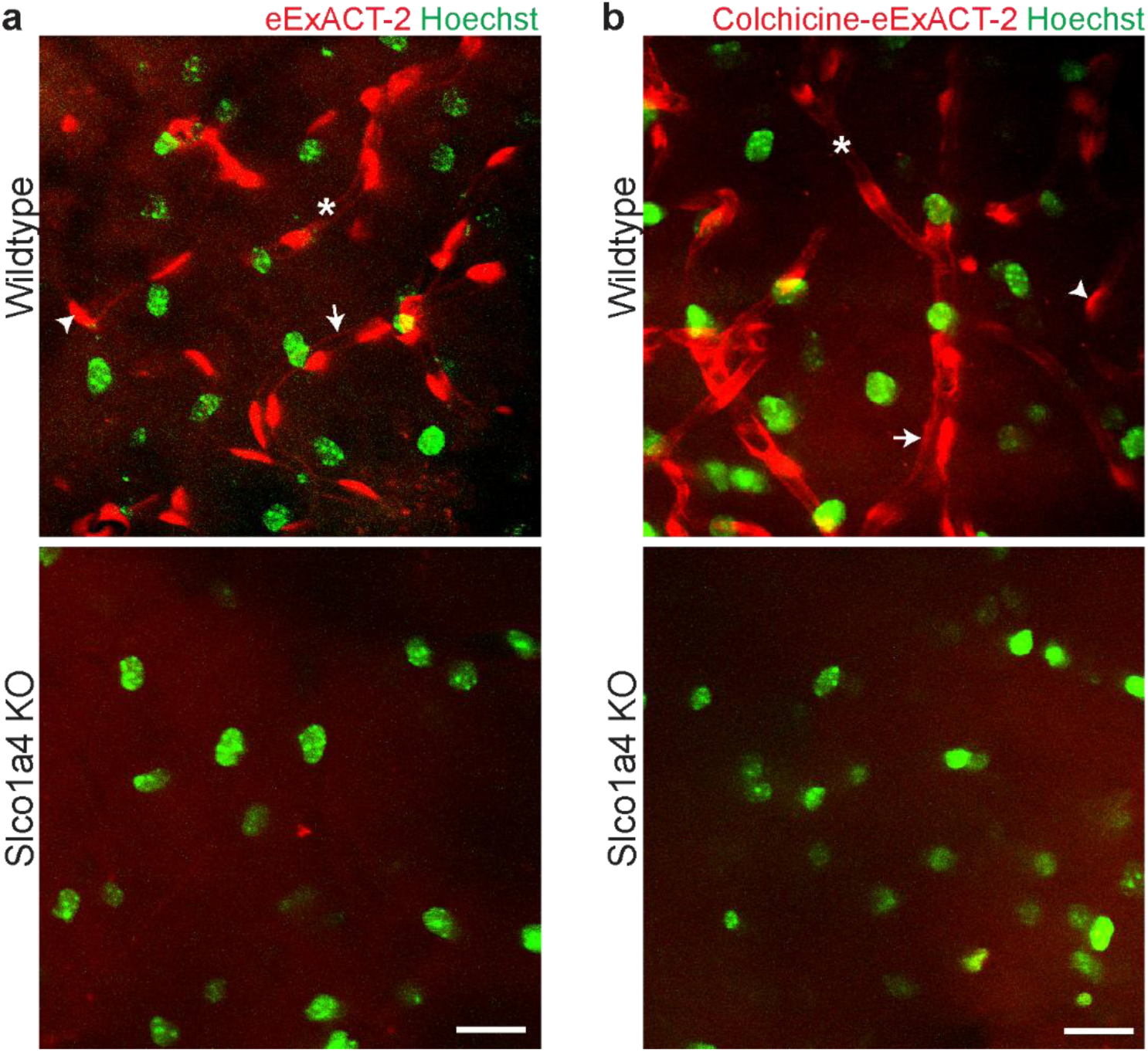
(Related to Figs. 2-3). The solute carrier Slco1a4 in mice facilitates the selective intracellular uptake of eExACT and colchicine-eExACT. **a.** *In vivo* two-photon images of the cortex from wild-type mice (top) and Slco1a4-deficient mice (bottom) after topical application of eExACT-2 (50 μM, red) and nuclear labeling with Hoechst (green). In wild-type mice, eExACT-2 labels endothelial processes (arrows) and cell bodies (arrowheads). Hoechst stains the nuclei in both wild-type and Slco1a4-deficient mice, providing additional contrast. In Slco1a4-deficient mice, the eExACT-2 signal is absent, confirming transporter dependence. Vessel lumen (asterisks). Scale bar, 20 μm. **b.** *In vivo* two-photon images of the cortex from wild-type mice (top) and Slco1a4-deficient mice (bottom) after topical application of Colchicine-eExACT-2 (50 μM, red) and nuclear labeling with Hoechst (green). In wild-type mice, Colchicine-eExACT-2 labels endothelial processes (arrows) and cell bodies (arrowheads). Hoechst stains the nuclei in both wild-type and Slco1a4-deficient mice, providing additional contrast. In Slco1a4-deficient mice, the Colchicine-eExACT-2 signal is absent, confirming transporter dependence. Vessel lumen (asterisks). Scale bar, 20 μm.

**Supplementary Figure 7.**
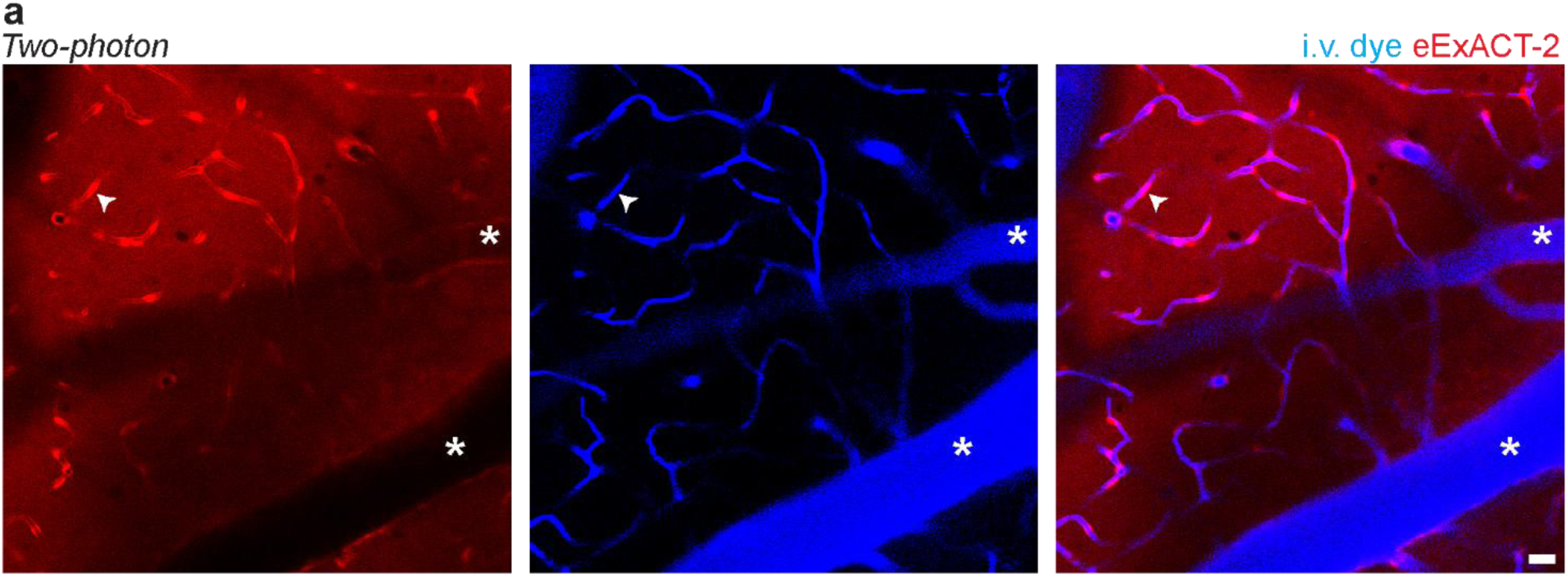
(Related to Figs. 1-2). eExACT selectively labels parenchymal microvasculature but not large leptomeningeal vessels. **a.** Two-photon images of the mouse cortex following topical application of eExACT-2 (red, 50 µM) and intravenous injection of a fluorescent vascular dye (blue). Large leptomeningeal surface vessels (asterisks) show strong intravascular dye signal but minimal eExACT labeling. In contrast, parenchymal microvessels exhibit robust endothelial uptake of eExACT-2 (arrowhead). Merged image (right) illustrates the selective labeling pattern. Scale bar, 20 μm.

**Supplementary Figure 8.**
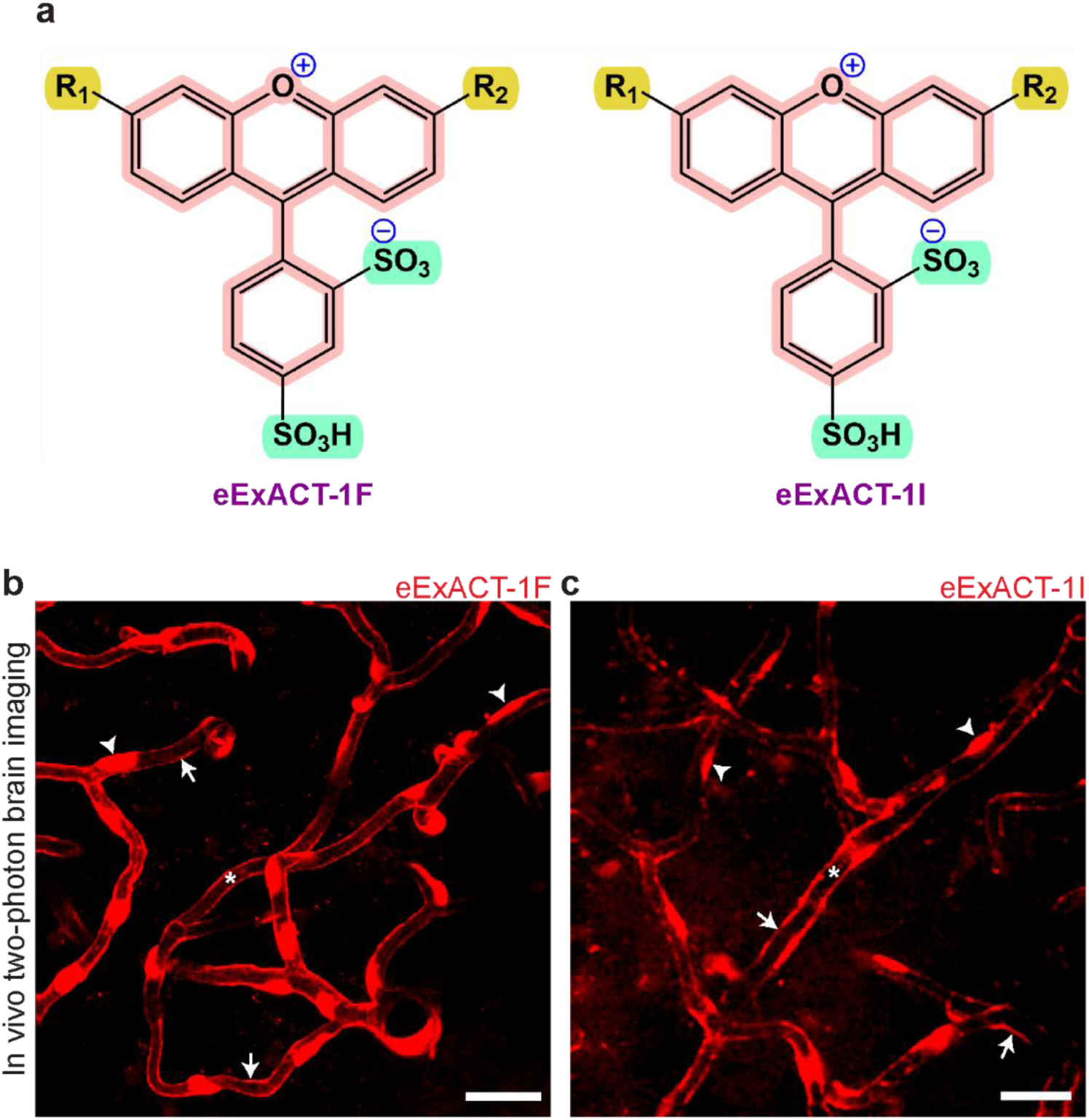
(Related to Fig. 3). Functional group modification with iodination or fluorination preserves endothelial targeting of eExACT. **a.** Chemical structures of fluorinated (eExACT-1F) and iodinated (eExACT-1I) derivatives of eExACT-1. **b–c.** Intravital two-photon images of mouse cortex following topical application of eExACT-1F (b) or eExACT-1I (c) (both at 50 μM) show robust and selective uptake by endothelial cells, including cell bodies (arrowheads) and processes (arrows), with minimal labeling of surrounding tissue. Asterisks denote vessel lumens. Scale bars, 20 μm.

**Supplementary Figure 9.**
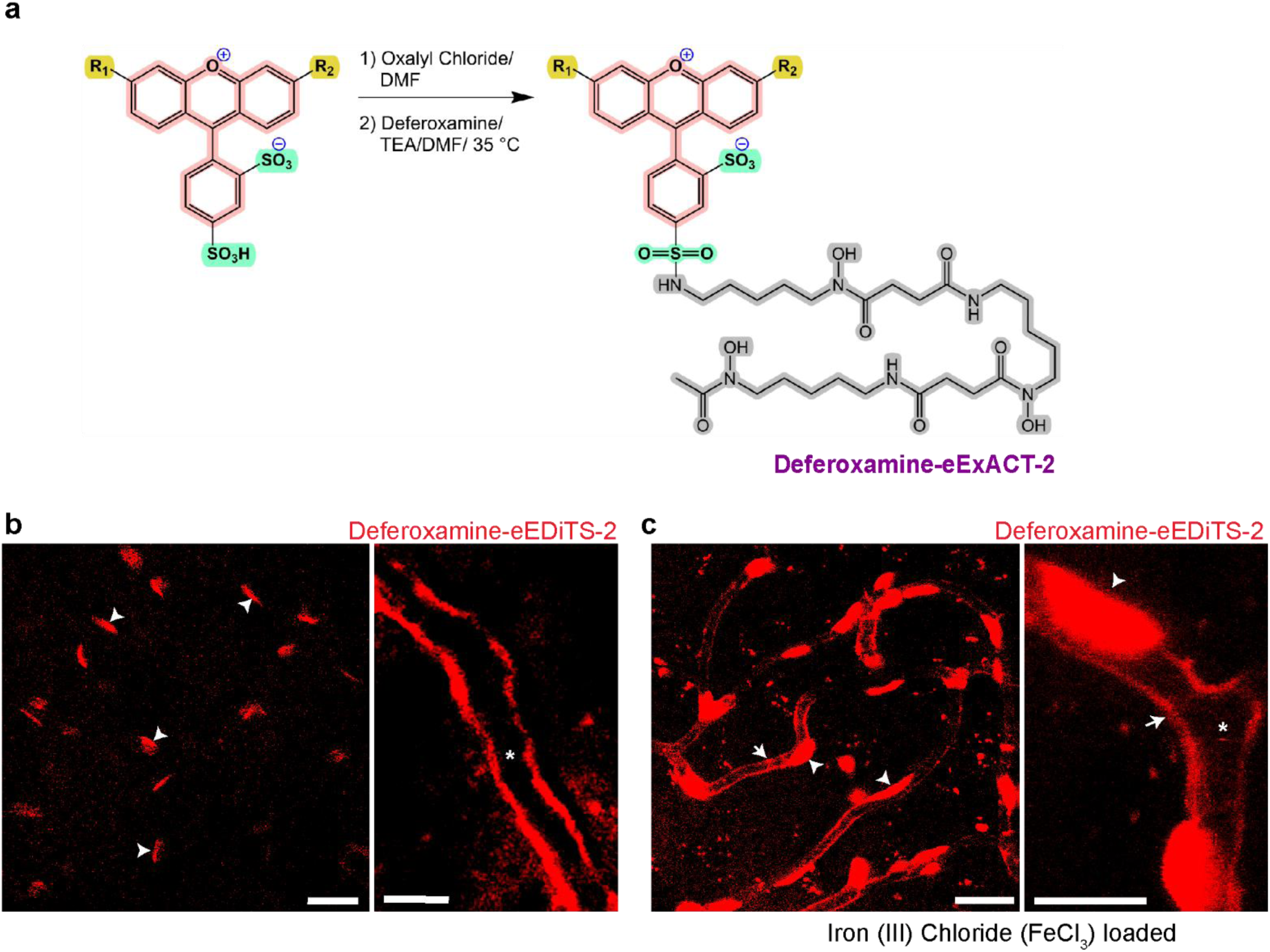
(Related to Fig. 3). eExACT conjugation to metal chelator deferoxamine preserves endothelial specificity even with trivalent iron encapsulation. **a.** Schematic representation of the synthetic route to generate Deferoxamine-eExACT-2. eExACT-2 was first activated with oxalyl chloride to form the acyl chloride intermediate, followed by conjugation with deferoxamine. **b.** Intravital two-photon images of mouse cortex following topical application of Deferoxamine-eExACT-2 (50 μM) show robust and selective uptake by endothelial cells, including cell bodies (arrowheads) with minimal labeling of surrounding cells. High-magnification view (right) showing Deferoxamine-eExACT-2 (50 μM, red) signals. Asterisks denote vessel lumens. Scale bars, 20 μm (left) and 10 μm (right). **c.** Intravital two-photon images of mouse cortex following topical application of Deferoxamine-eExACT-2 (50 μM) pre-treated with 200 µM FeCl₃ show robust and selective uptake by endothelial cells, including cell bodies (arrowheads) and processes (arrows). Asterisks denote vessel lumens. High-magnification view (right) showing Deferoxamine-eExACT-2 (50 μM, red) signals. Cell bodies (arrowheads); processes (arrows), and vessel lumen (asterisks). Scale bars, 20 μm (left) and 10 μm (right).

**Supplementary Figure 10.**
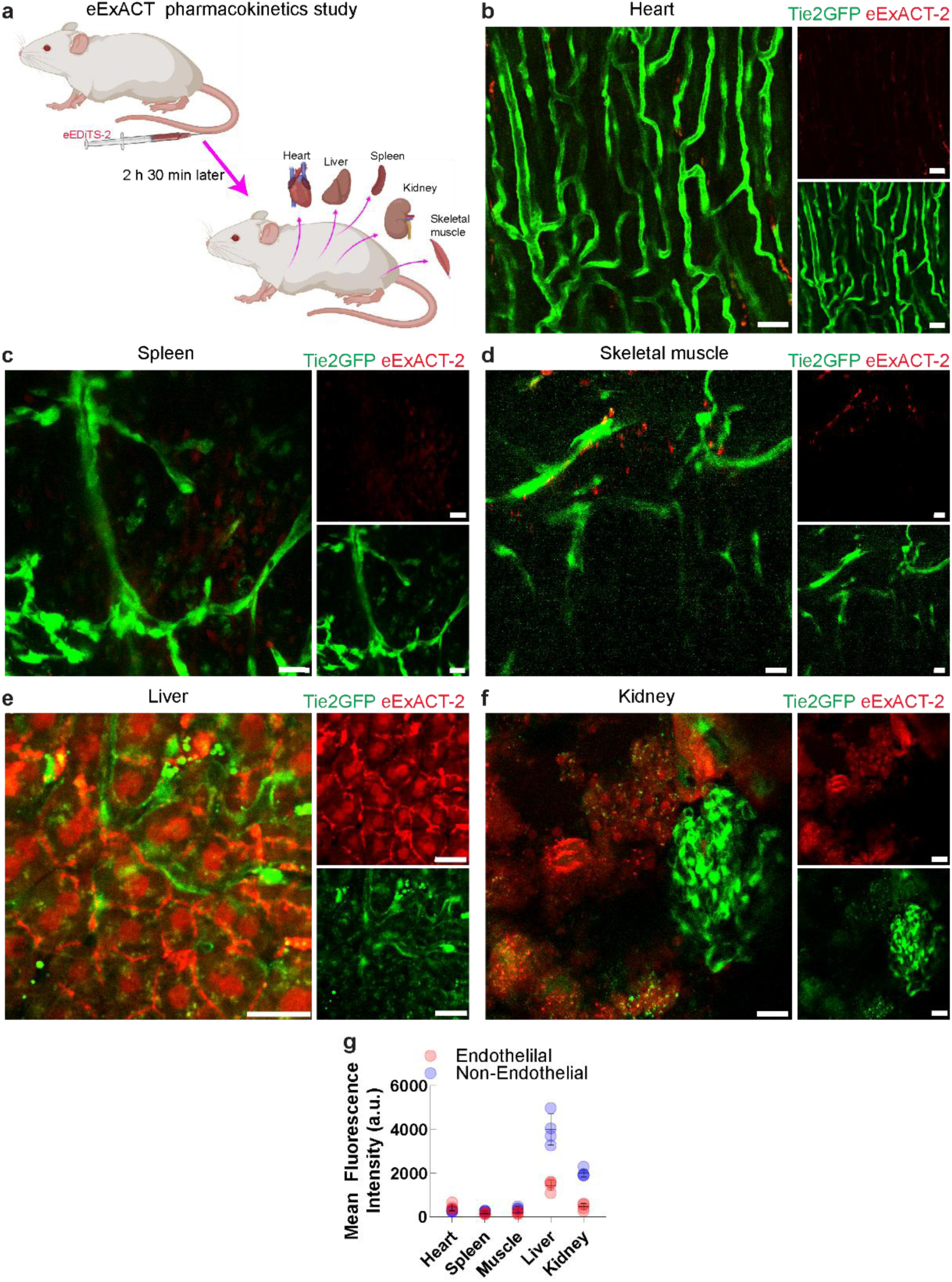
(Related to Fig. 5). Biodistribution of eExACT in various organs following intravenous administration. **a.** Schematic of the pharmacokinetic experiment. Mice received an intravenous injection of eExACT-2 (0.1 mM, 70 µL) and were sacrificed 2.5 hours later for imaging of heart, spleen, skeletal muscle, liver, and kidney. **b–f**. Two-photon images of explanted organs from Tie2-GFP mice showing endothelial cells (green) and eExACT-2 signal (red) in heart (b), spleen (c), skeletal muscle (d), liver (e), and kidney (f). Liver hepatocytes and kidney tubular epithelial cells show prominent uptake of eExACT-2. In contrast, endothelial cells in all peripheral organs, including those lining the liver sinusoids and the glomerular capillaries in the kidney, display minimal to no uptake. **g.** Other parenchymal cell types in heart, spleen, and skeletal muscle also exhibit negligible labeling compared to liver and kidney. This distribution pattern highlights the strong selectivity of eExACT-2 for brain endothelium. Insets show separated and merged fluorescence channels. Scale bars, 20 μm.

**Supplementary Figure 11.**
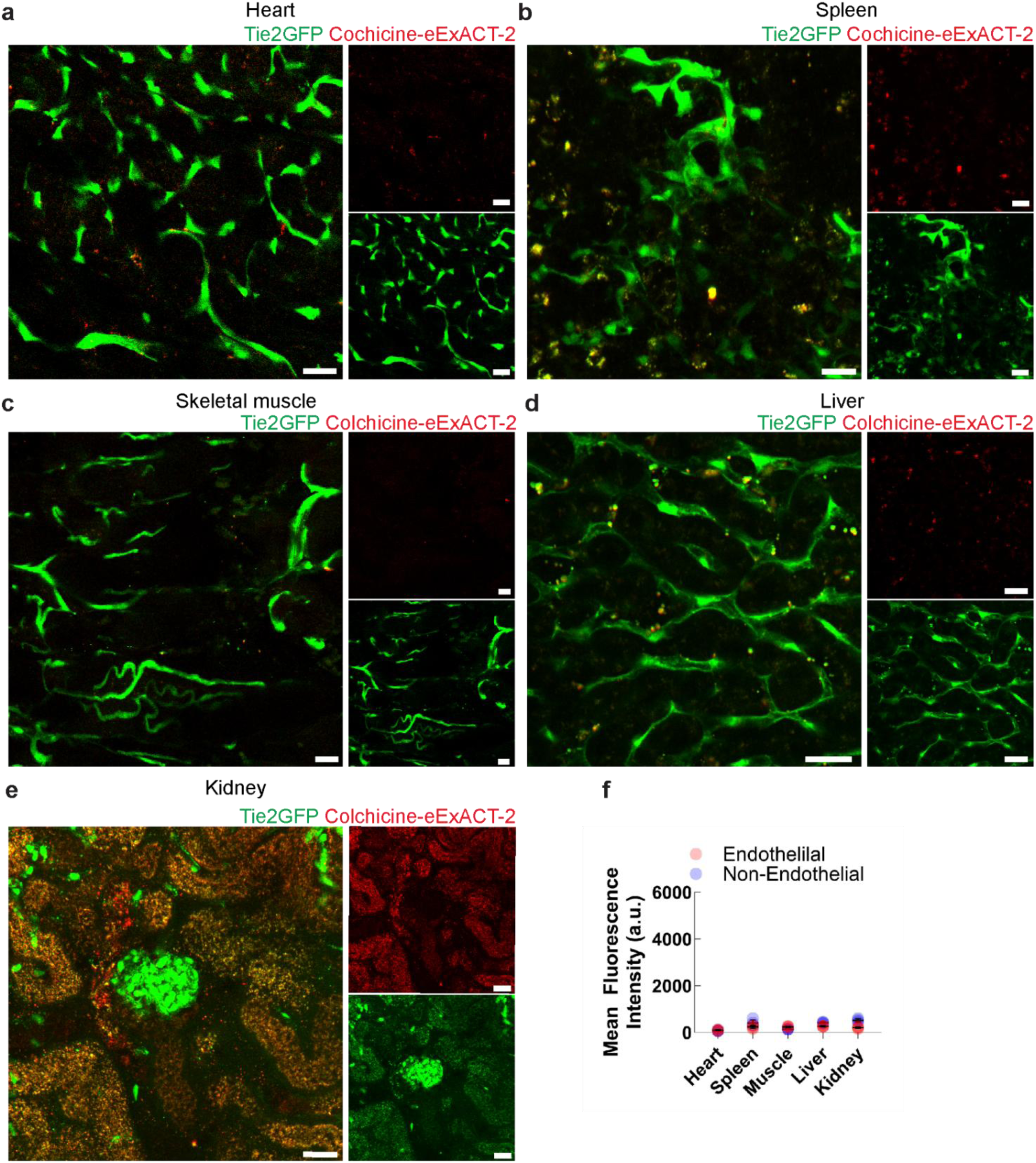
(Related to Fig. 5). Biodistribution of Colchicine-eExACT in various organs following intravenous administration. Mice received an intravenous injection of Colchicine-eExACT-2 (0.1mM, 70 µL) and were sacrificed 2.5 hours later for imaging of heart, spleen, skeletal muscle, liver, and kidney. **a–e**. Two-photon images of explanted organs from Tie2-GFP mice showing endothelial cells (green) and eExACT-2 signal (red) in heart (a), spleen (b), skeletal muscle (c), liver (d), and kidney (e). Liver hepatocytes and kidney tubular epithelial cells show low levels of uptake of colchicine-eExACT-2. In contrast, endothelial cells in all peripheral organs, including those lining the liver sinusoids and the glomerular capillaries in the kidney, display minimal to no uptake. Other parenchymal cell types in heart, spleen, and skeletal muscle also exhibit negligible labeling. **f.** Uptake quantification shows overall low levels compared to unconjugated eExACT-2 (see Supplementary Figure 10g). Insets show separated and merged fluorescence channels. Scale bars, 20 μm. Note: Mice received an intravenous injection of Colchicine-eExACT-2 and were sacrificed 2.5 hours later for imaging of heart, spleen, skeletal muscle, liver, and kidney.

**Supplementary Figure 12.**
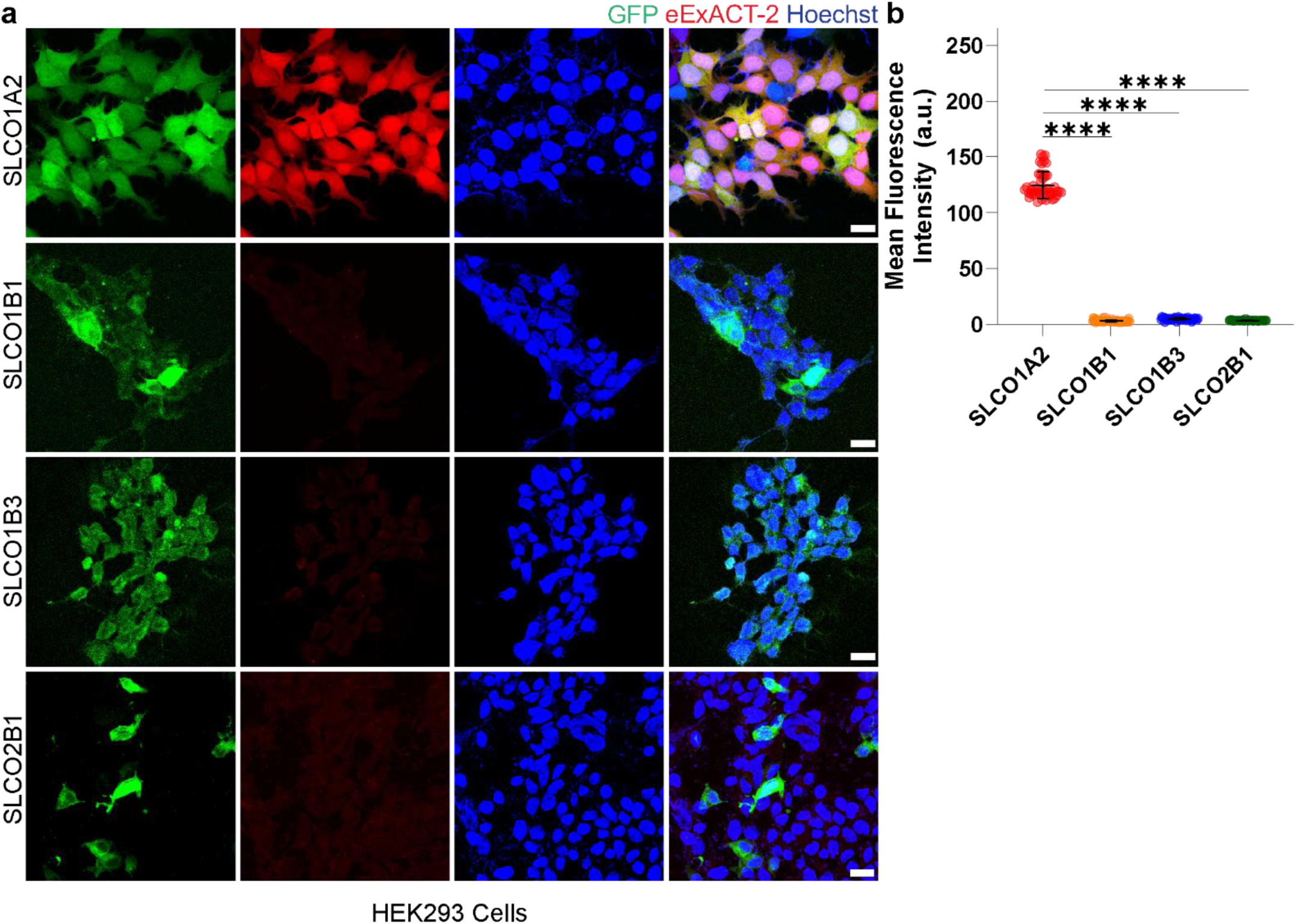
(Related to Fig. 5). eExACT-2 is not taken up by cells expressing SLCO1B1, SLCO1B3, or SLCO2B1. **a.** Representative confocal fluorescence microscopy images of HEK293 cells expressing SLCO1A2, SLCO1B1, SLCO1B3, or SLCO2B1, incubated with eExACT-2 (red, 10 μM). Columns show fluorescence from GFP (green, indicating transporter expression), eExACT-2 (red), and nuclei stained with Hoechst (blue). The merged images demonstrate strong co-localization of eExACT-2 with SLCO1A2-expressing cells, while minimal to no uptake is observed in cells expressing SLCO1B1, SLCO1B3, or SLCO2B1. Scale bars: 20 µm. **b.** Quantification of mean eExACT-2 (10 μM) fluorescence intensity confirms a lack of probe accumulation in SLCO1B1-, SLCO1B3-, and SLCO2B1-expressing cells, in contrast to the strong uptake observed in SLCO1A2-expressing cells (related to Fig. a). Data are presented as mean ± SD (****p < 0.0001, Two-tailed unpaired t-test with Welch’s correction).

**Supplementary Figure 13.**
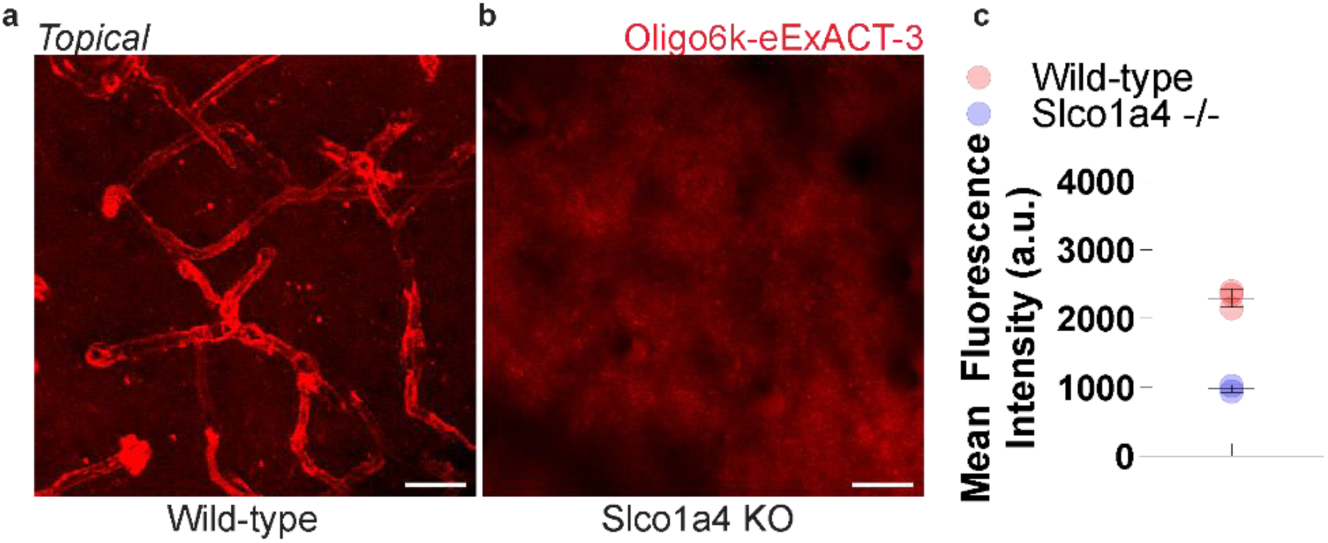
(Related to Fig. 6). SLCO1A4-Dependent in Vivo Endothelial Uptake of High-Molecular-Weight Oligonucleotides via eExACT Conjugation. **a-b.** Representative intravital two-photon images of the cortex in (a) wild-type and (b) Slco1a4 knockout mice following topical application of 6000 Da oligonucleotide-conjugated eExACT-3 (Oligo6k-eExACT-3, 60 µM). Robust in vivo uptake into brain endothelial cells is observed in wild-type mice (a), while uptake is abolished in Slco1a4 knockout mice (b), demonstrating transporter-dependent endothelial uptake. Scale bars, 20 µm. **c.** Quantification of mean fluorescence intensity in brain endothelial cells of wild-type versus Slco1a4-deficient mice, measured 30 minutes post-cortical application of Oligo6k-eExACT-3 (60 µM).

**Supplementary Figure 14.**
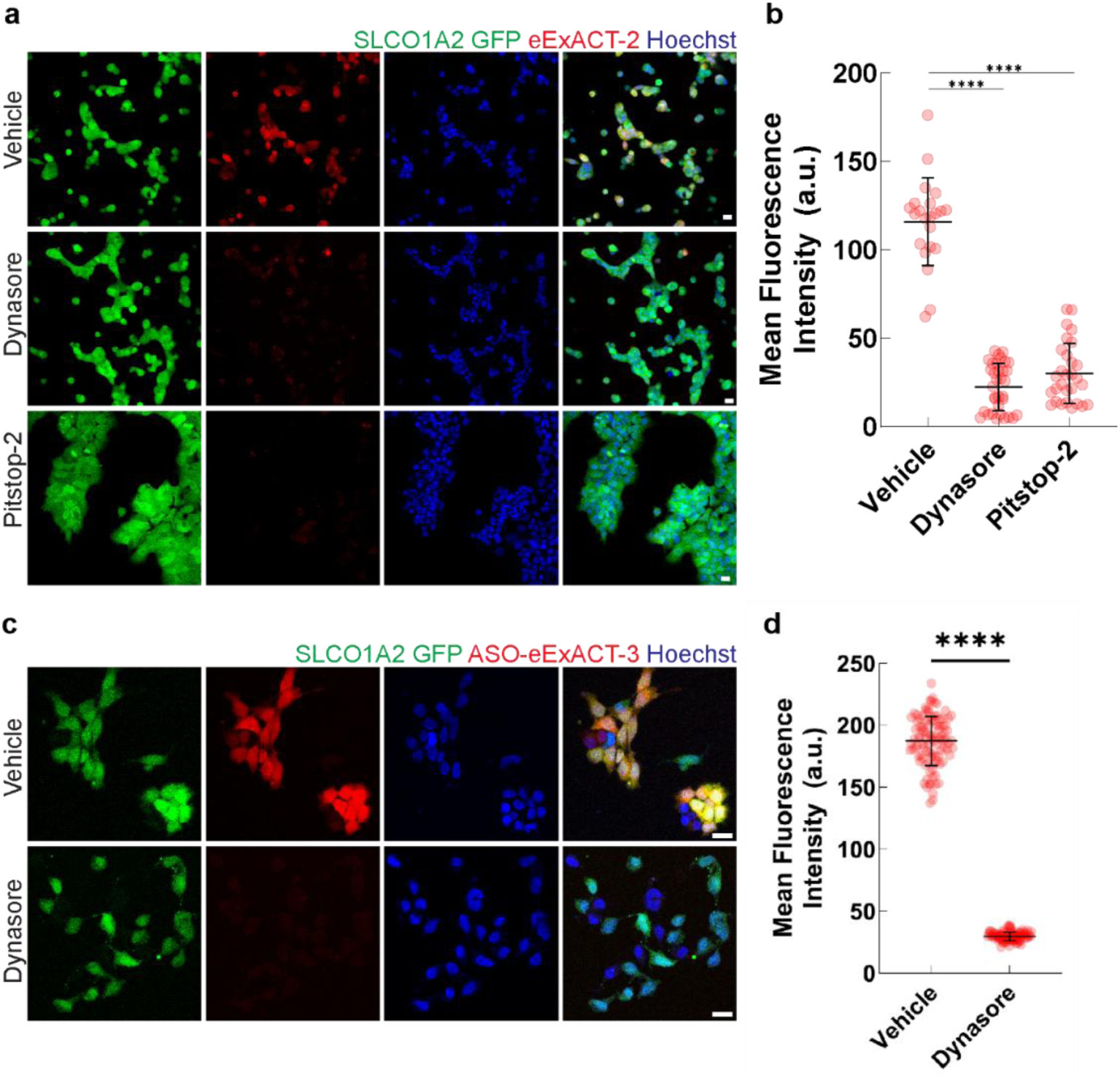
(Related to Fig. 6). Endocytosis contributes to SLCO1A2-dependent uptake of eExACT conjugates. **a.** Confocal fluorescence images of SLCO1A2-expressing HEK293 cells treated with eExACT-2 in the presence of pitstop-2, dynasore, or vehicle. GFP (green, SLCO1A2), eExACT (red), Hoechst (blue). Vehicle-treated cells show strong eExACT-2 colocalization with SLCO1A2; uptake is abolished by pitstop-2 or dynasore, supporting an endocytosis-dependent mechanism. Scale bars, 20 µm. All compounds at 10 µM unless noted. **b.** Quantification of mean fluorescence intensity for eExACT-2 confirms absence of probe accumulation in pitstop-2- or dynasore-treated cells versus vehicle-treated SLCO1A2-expressing cells. Mean ± SD; ****p < 0.0001, unpaired t-test with Welch’s correction. **c.** Confocal fluorescence images of SLCO1A2-expressing HEK293 cells treated with ASO-eExACT-3 with dynasore or vehicle. GFP (green), ASO-eExACT-3 (red), Hoechst (blue). Dynasore markedly reduces probe uptake relative to vehicle, consistent with endocytosis-dependent SLCO1A2-mediated internalization. Scale bars, 20 µm. All compounds at 10 µM unless noted. **d.** Quantification of mean fluorescence intensity for ASO-eExACT-3 confirms significantly reduced accumulation in dynasore-treated versus vehicle-treated SLCO1A2-expressing cells. Mean ± SD; ****p < 0.0001, unpaired t-test with Welch’s correction.

**Supplementary Figure 15.**
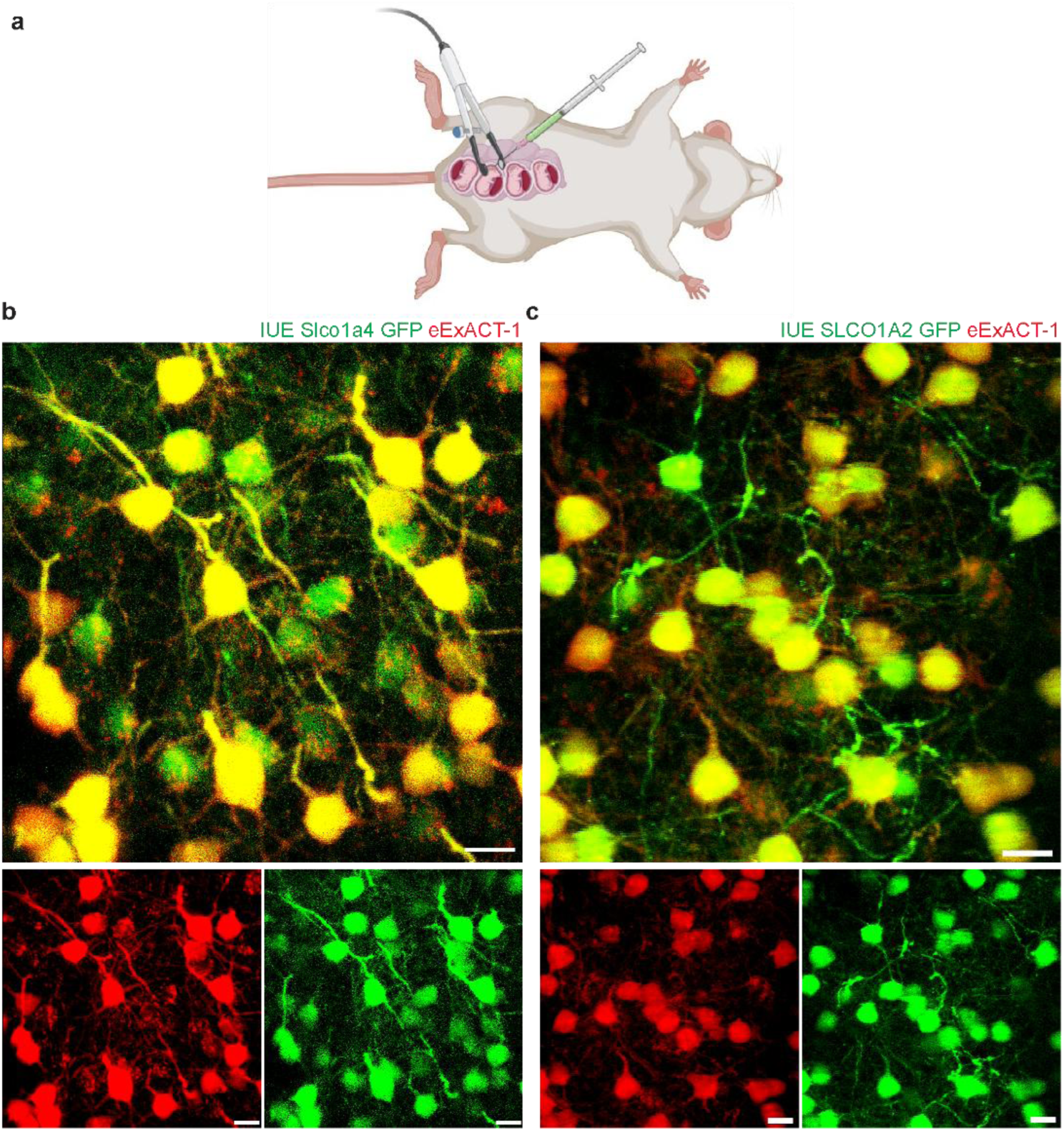
(Related to Fig. 7). Transfection of cortical neurons via in utero electroporation of either Slco1a4 or SLCO1A2 leads to robust uptake of eExACT in neurons. **a.** Schematic illustration of the in-utero electroporation (IUE) procedure used to introduce transporter constructs and a GFP reporter into the developing cortex. **b-c.** *In vivo* two-photon images showing selective uptake of eExACT-1 (red, 50 µM) in neurons co-expressing GFP (green) and either Slco1a4 (b, left) or SLCO1A2 (c, right). Insets show individual fluorescence channels. Strong dye uptake is observed throughout neuronal cell bodies and processes, including dendrites. Notably, endothelial uptake is reduced in regions of neuronal expression, suggesting competitive uptake of the compound. Scale bars, 10 μm.

**Supplementary Figure 16:**
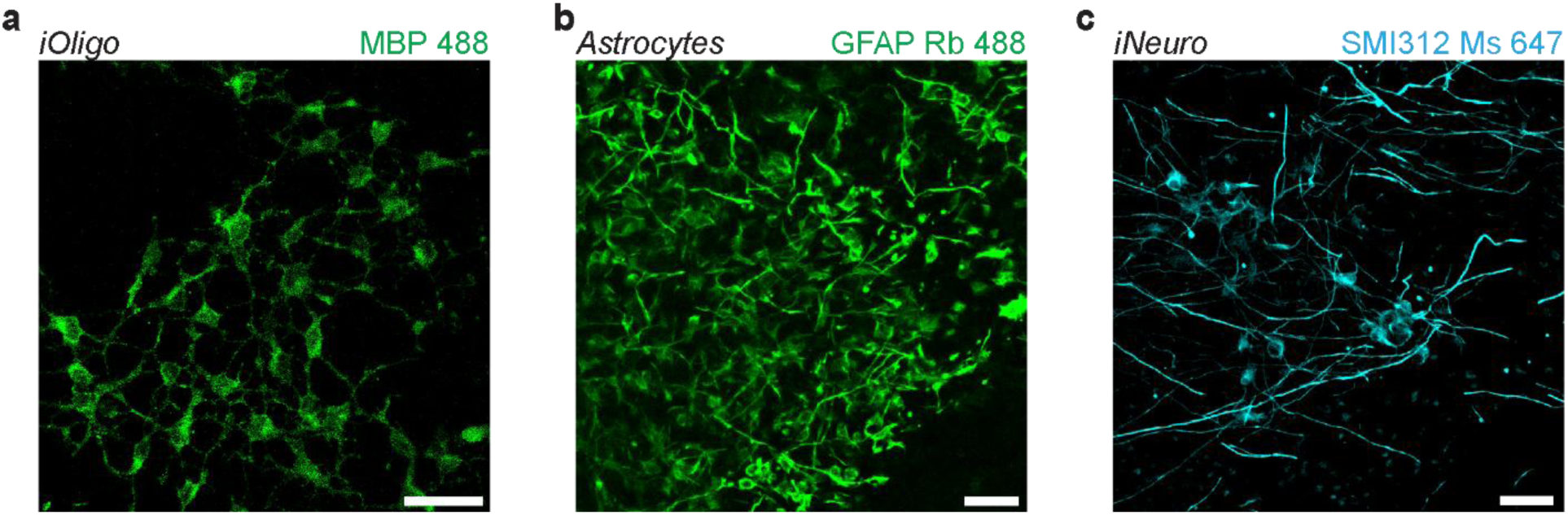
(Related to Fig. 7). Validation of cell identity for iPSC-derived and human primary cell types by immunofluorescence characterization. **a.** iPSC-derived oligodendrocytes (iOligo) immunostained for Myelin Basic Protein (MBP; rabbit, Alexa Fluor 488, green), demonstrating characteristic morphology and myelination-associated processes. **b.** Human astrocytes immunolabeled with anti-GFAP (rabbit, Alexa Fluor 488, green), revealing the stellate, highly ramified morphology typical of mature astrocytes. **c.** iPSC-derived neurons (iNeuro) stained with SMI312, a pan-axonal neurofilament marker (mouse, Alexa Fluor 647, cyan), highlighting neuronal soma and axonal/dendritic projections. Expression of lineage-specific markers confirms cell identity prior to downstream experiments. All images were acquired by confocal microscopy. Scale bars, 50 µm.

**Supplementary Figure 17.**
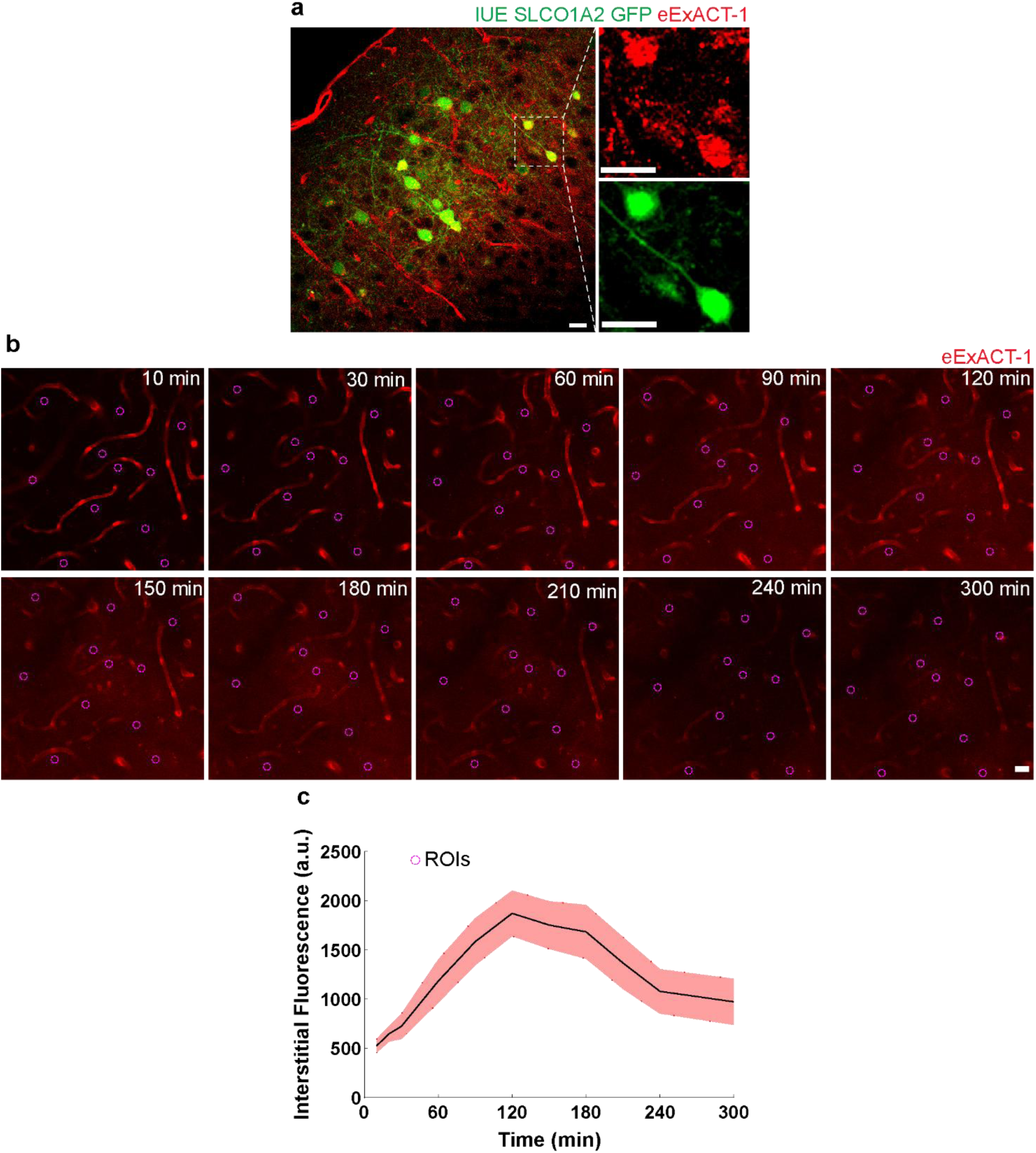
(Related to Fig. 7). eExACT crosses the blood–brain barrier and accumulates in the brain parenchyma. **a.** Confocal image of fixed cortical tissue from a mouse previously transfected with SLCO1A2 via in utero electroporation, following intravenous injection of eExACT-1 (100 µL of 0.1 mM). Robust neuronal labeling indicates successful BBB penetration and transporter-mediated uptake. Scale bars, 20 μm. **b.** In a separate experiment, eExACT-1 (70 µL of 0.1 mM, red) was intravenously injected, and *in vivo* two-photon microscopy was performed through a cranial window. Fluorescence was measured in extravascular regions (dotted ROIs) across multiple fields of view. **c.** Quantification of fluorescence intensity over time reveals progressive accumulation of eExACT-1 outside the vasculature, consistent with effective BBB transit. n = 20 fields per mouse, 2 mice total.

